# Using single-cell models to predict the functionality of synthetic circuits at the population scale

**DOI:** 10.1101/2021.08.03.454887

**Authors:** Chetan Aditya, François Bertaux, Gregory Batt, Jakob Ruess

**Affiliations:** Inria Paris, 2 rue Simone Iff, 75012 Paris, France; Institut Pasteur, 28 rue du Docteur Roux, 75015 Paris, France; Université de Paris, 85 Boulevard Saint-Germain, 75006 Paris, France

**Author notes:** contributed equally (Gregory Batt, Jakob Ruess).

**Keywords:** Optogenetics, synthetic differentiation circuits, composability, chemical master equation, population dynamics

## Abstract

Mathematical modeling has become a major tool to guide the characterization and synthetic construction of cellular processes. However, models typically lose their capacity to explain or predict experimental outcomes as soon as any, even minor, modification of the studied system or its operating conditions is implemented. This limits our capacity to fully comprehend the functioning of natural biological processes and is a major roadblock for the de novo design of complex synthetic circuits. Here, using a specifically constructed yeast optogenetic differentiation system as an example, we show that a simple deterministic model can explain system dynamics in given conditions but loses validity when modifications to the system are made. On the other hand, deploying theory from stochastic chemical kinetics and developing models of the system’s components that simultaneously track single-cell and population processes allows us to quantitatively predict emerging dynamics of the system without any adjustment of model parameters. We conclude that carefully characterizing the dynamics of cell-to-cell variability using appropriate modeling theory may allow one to unravel the complex interplay of stochastic single-cell and population processes and to predict the functionality of composed synthetic circuits in growing populations before the circuit is constructed.

## Introduction

At the heart of rational circuit design in synthetic biology lies the assumption that the functionality of complex circuits can be predicted from known properties of their components. Yet in practice, we routinely fail to make predictions of circuit dynamics that would agree with the data at the level expected in physics or engineering. A core reason for this is cell-to-cell variability inside genetically identical cell populations. Such population heterogeneity is often a consequence of the inherent stochasticity of biochemical processes inside cells [1]. Cell-to-cell variability may lead to unexpected and undesirable circuit dynamics and has been identified as one of the major roadblocks for designing synthetic circuits with in-silico predictable functionality [2, 3]. On the other hand, it has been shown that identifying and carefully characterizing sources of variability at the single-cell scale allows one to design remarkably robust synthetic circuits [4] or to exploit stochasticity for creating features of cell populations, such as bimodal phenotype distributions, that would otherwise be difficult to engineer [5].

While the single-cell perspective has certainly helped to advance our understanding of cellular processes [6, 7, 8, 9], what eventually matters for most applications in synthetic biology is how a particular circuit functions inside growing populations. At the population-scale, synthetic circuits may intentionally (e.g. circuits to control growth [10]) or unintentionally (e.g. burden caused by protein production [11]) affect traits of cells that can be selected during population growth. If this is the case, we need to expect that variability that is generated at the single-cell scale (e.g. stochastic production of the burdensome protein) will lead to consequences at the population-scale that cannot be predicted solely based on a characterization of the circuit inside single cells. However, despite the apparent prevalence of problems where single-cell and population processes are coupled, multi-scale models that capture both single-cell stochastic chemical kinetics as well as population dynamics have not been used to predict, or even just to explain, how population-scale functionality emerges from single-cell characteristics of synthetic circuits. Presumably, this is because the classically used modeling framework for single-cell processes, that is the chemical master equation [12, 13], is only directly applicable at the population-scale whenever the population phenotype distribution is equivalent to the distribution of the single cell process and not additionally shaped by population-level processes [14].

Here, we construct an artificial yeast differentiation system in which a Cre-recombinase is expressed from a light-inducible promoter and used to create dynamically controllable yeast communities of differentiated and undifferentiated cell types [15]. The system is equipped with four fluorescent reporter proteins to allow for simultaneous measurements of cellular processes and the differentiation state of cells. The core feature of our system is that varying the duration of applied light pulses allows one to modulate the fraction of cells that differentiates even though the entire population is exposed to the same global light stimulation. The system therefore exploits heterogeneity in the response of cells to light to enable external control of population dynamics but, by doing so, creates a complex coupling between population dynamics and the single-cell process that leads to the production of recombinase. It can therefore serve as an ideal test-bed to study what types of models are needed to explain and predict complex dynamics of synthetic circuits at the population scale. We find that a simple deterministic model that ignores all sources of variability can explain dynamics of a chromosomally integrated system version fairly well. However, modifying sources of variability and growth conditions, by expressing the system from plasmids and growing cells in selective media, leads to structural incapacity of the deterministic model to match measured population dynamics. To remedy this shortcoming, we derive a non-linear version of the chemical master equation (CME) that tracks conditional probabilities and that can be used to simultaneously model single-cell and population processes. Subsequently, we develop one such non-linear CME model to represent the chromosomally integrated differentiation system and another to track plasmid copy number fluctuations, plasmid loss, and growth in selective media. We find that composing these two models allows one to not only match, but to in-silico predict, complex emerging dynamics of the plasmid-based differentiation system without any modification of model parameters. These results suggest that the frequently encountered failure of model predictions for composed synthetic circuits may be resolvable with appropriate component models that are carefully constructed to track the coupled dynamics of single-cell and population processes via multi-scale stochastic kinetic models.

### A yeast optogenetic differentiation system

To be able to study the role of cell-to-cell variability in the design and functionality of synthetic circuits, we constructed a yeast optogenetic differentiation system whose dynamics can be well-observed at the single-cell level thanks to the simultaneous presence of four different fluorescent reporters. Concretely, we started from a circuit design that we recently published [15] and used a Cre-recombinase (Cre) under control of the EL222 promoter [16] to trigger differentiation in controllable population fractions using global light stimulation patterns (Figure 1). When expressed, Cre excises a DNA fragment that is designed such that, upon recombination, cells switch from constitutively producing blue fluorescent protein (mCerulean) to producing green fluorescent protein (mNeonGreen). Furthermore, red fluorescent protein (mScarlet-I) is produced alongside Cre from a second copy of the EL222 promoter and provides an observable readout that reports on Cre expression levels in response to light. Finally, EL222 transcription factor (EL222) is constitutively expressed from a pTDH3 promoter and fused to a yellow fluorescent protein (mVenus). In summary, the system allows one to trigger and monitor recombination in cells and to observe correlations of the probability for cells to recombine with cellular amounts of EL222. To test system functionality, we performed experiments using a platform of LED-equipped and fully computer-controlled parallel bioreactors [17]. Single-cell measurements are automatically taken by flow cytometry with the help of a programmable pipetting robot (Figure 1b). Using deconvolution to extract amounts of the different reporter proteins in cells from measured spectral signatures (see Supporting Information, Section S.7), we find that all cells gradually switch from blue to green when sufficient light is applied. On the other hand, exposing cell populations to pulses of light leads to bimodal mNeonGreen distributions (Figure 1c and Supporting Information, Figure S.1), which shows that only a fraction of the population recombines in response to light pulses. Applying a threshold in mNeonGreen fluorescence to classify cells into recombined and non-recombined, we can quantify the differentiation dynamics of the system (Figure 1c,d). We find that the system’s response to light can be captured fairly well by a simplistic population dynamics model that relates differentiated and undifferentiated cells via a constant differentiation rate in the presence of light (Figure 1d and Supporting Information, Section S.1). To test if the probability for a cell to recombine is correlated with single-cell levels of EL222 and Cre, we analyzed mVenus and mScarlet-I fluorescence distributions shortly after applied light pulses. We find only minor differences in EL222 levels of undifferentiated cells before and after light induction that are difficult to distinguish from small inaccuracies in deconvolution or reactor-to-reactor variability of the experimental platform (Figure 1c and Figure S.1). Overall, we are led to conclude that cell-to-cell variability in EL222 and Cre can be safely ignored and that the functionality of the system can readily be characterized by a simple deterministic model. However, past experience in synthetic biology has shown that most circuits only function reliably in tightly constrained operating conditions and even seemingly good models retain their predictive power only in the precise context that has been used to construct the model.

**Figure 1:**
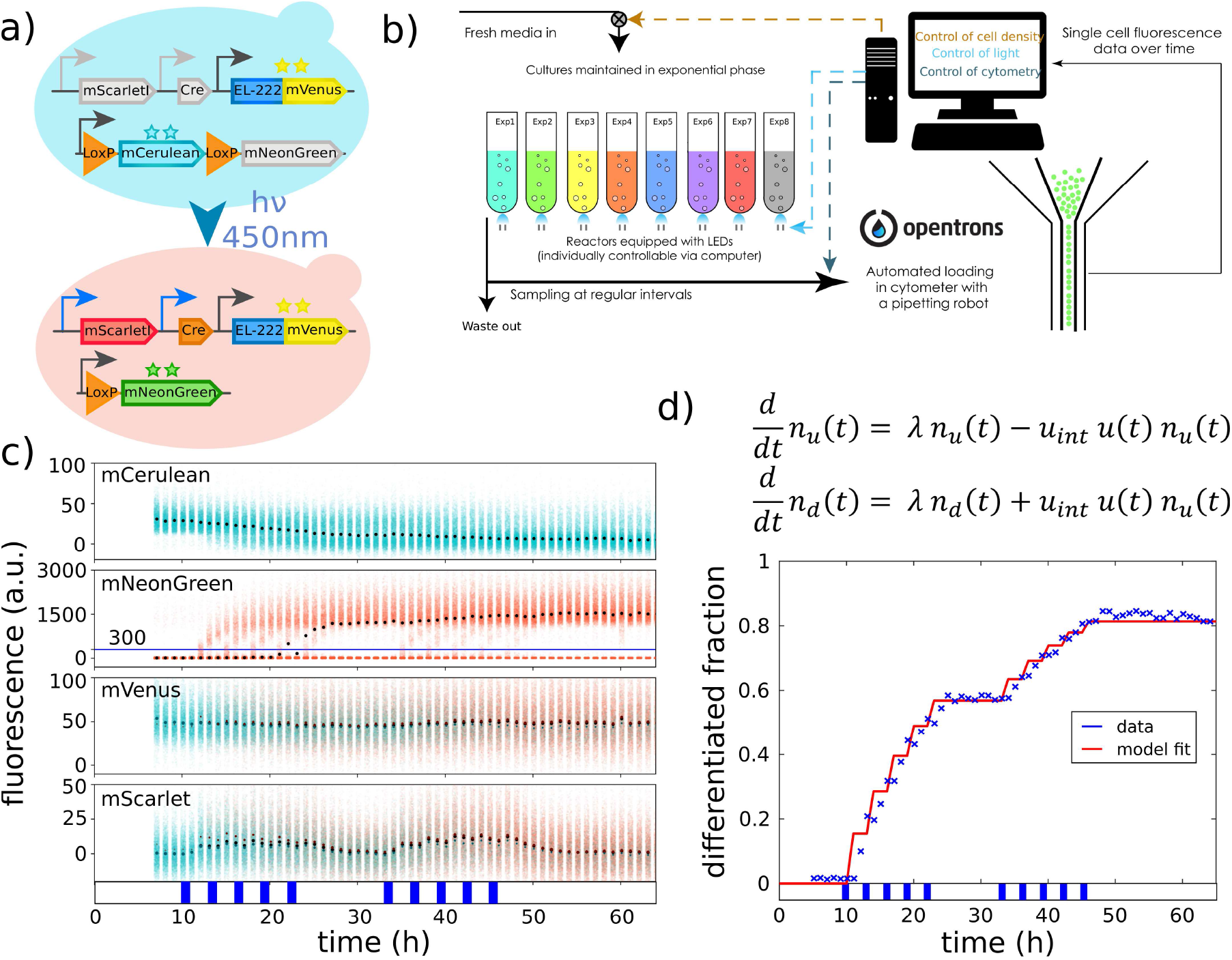
A yeast optogenetic differentiation system. **(a)** EL222 is produced in cells from a constitutive promoter (pTDH3, dark grey). Upon exposure of cells to blue light, EL222 switches the EL222-promoter from off (light grey) to on (blue) and cells start to produce mScarlet-I and Cre-recombinase. Cre excises a DNA-fragment placed between two target LoxP-sites which is designed such that cells switch from producing mCerulean to mNeonGreen from a second constitutive promoter (pTDH3, dark grey). **(b)** Yeast cells harboring the differentiation system are grown in parallel, fully automated bioreactors that are equipped with controllable blue LEDs. Optical density of cultures is maintained constant and flow cytometry measurements are taken at regular intervals via a custom-programmed pipetting robot. **(c)** Upon exposure to light, mScarlet-I fluorscence of some cells increases quickly, which indicates that also Cre is produced. Accordingly, mCerulean fluorescence decreases and mNeonGreen fluorescence increases. mNeonGreen fluorescence is then used to classify cells into differentiated and un-differentiated (blue line). EL222, fused to mVenus, is constitutively expressed in all cells and not affected by recombination. Small changes in mVenus levels in response to light that are noticeable in the data (smeared out, wider distributions towards the end) are due the presence of mNeonGreen in cells, which leads to small errors in the deconvolution of mVenus fluorescence (Supporting Information, Section S.7). Dots show median fluorescence in all panels (black: all cells, cyan: undifferentiated cells, orange: differentiated cells). The applied light sequence is shown at the bottom (two times five 1h pulses with 2h between subsequent pulses). The data is also visualized in the form of distributions in the Supporting Information, Figure S.1. **(d)** A simple model (top) is capable of capturing the population dynamics of differentiated and undifferentiated cells very well. The applied light sequence is shown at the bottom.

### Modifying circuits leads to unpredictable functionality

To test if the functionality of our differentiation system remains predictable when the circuit is modified, we constructed a variant of the system in which EL222 and Cre genes are placed on 2-micron plasmids instead of being chromosomally integrated (Figure 2a).

**Figure 2:**
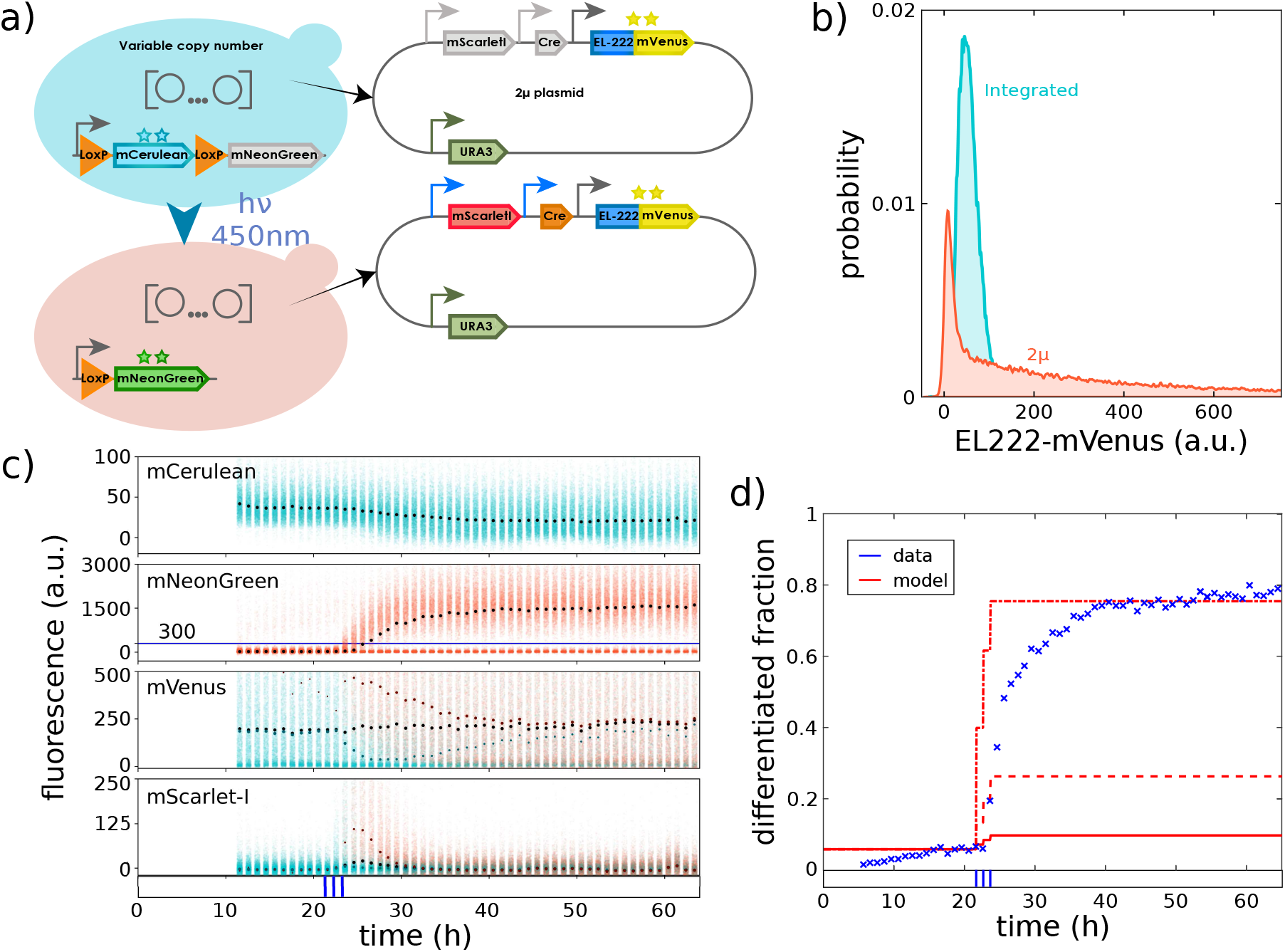
Expressing the differentiation system from plasmids breaks straightforward predictability. **(a)** To analyze the consequences of changes in cell-to-cell variability, a new strain was constructed in which Cre and EL222 genes are placed on 2-micron plasmids instead of being chromosomally integrated. **(b)** Measuring stationary EL222 distributions in the dark for the plasmid-based version of the system (red) shows that cell-to-cell variability in the amount of EL222 is strongly increased compared to the integrated version of the system (blue). **(c)** Large cell-to-cell variability in EL222 propagates to large variability in mScarlet-I (and thus Cre) production with some cells displaying much higher mScarlet-I fluorescence compared to the integrated system version (5-fold increased scale on y-axis for mScarlet-I and mVenus compared to Figure 1c). Using mNeonGreen fluoresence to classify cells into differentiated and undifferentiated (blue line) shows that light causes a transient split in mVenus distributions of the two sub-populations in response to light. The applied light sequence is shown at the bottom (three 5min pulses with 55min between subsequent pulses). **(d)** The simple model of Figure 1d fails to capture population dynamics for the plasmid-based version of the system. *Crosses:* measured differentiated population fractions. *Solid line:* model predictions using the same parameter values as in Figure 1d for the integrated system. *Dashed line:* model predictions after adjusting the differentiation rate parameter to account for the presence of multiple copies of the system. *Dashdotted line:* re-fitting the model to match final stationary differentiated fractions (30-40h after first light induction) leads to significant model mismatch during early transient dynamics. The applied light sequence is shown at the bottom.

Since plasmid copy numbers vary between cells, we expect this change to lead to significant differences in EL222 and Cre average levels and additional cell-to-cell variability. Indeed, growing cells in the dark, we find that EL222 distributions are very different from the integrated system version and characterized by much heavier tails and a mode that is shifted to lower levels (Figure 2b). Taken together, these two features imply that on average cells in the population contain more EL222 (almost 6-fold) but at the same time more cells contain less EL222 compared to the integrated system version (which notably falsifies the a priori expectation that all cells would have higher EL222 levels due to the presence of multiple plasmids in cells). We may therefore wonder if and how these differences impact the functionality of the circuit and if emerging dynamics of the population composition can still be predicted by the simple model in Figure 1d. Exposing cells to different light patterns, we find that the same amount of light leads to differentiation of more cells for the plasmid based version of the system (Figure S.3), which is in line with higher average levels of EL222 in the population and the presumable presence of multiple plasmids in cells, each carrying a copy of the promoter driving Cre. Adjusting the differentiation rate parameter in the simple deterministic model to account for the on-average presence of multiple copies of the system, however, does not lead to agreement of model predictions and data, nor is it at all possible to obtain any precise fit of this model to the population dynamics that emerge from the plasmid-based differentiation system (Figure 2d). Analysing mVenus distributions, we find that differentiated cells shortly after applied light pulses are characterized by high EL222 levels whereas the mVenus distribution of the undifferentiated subpopulation is shifted to lower levels compared to the total mVenus population distribution (Figure 2c). Since EL222 is constitutively expressed from the same promoter and plasmids in all cells, we conclude that these differences must be caused by selective differentiation of cells with high amounts of EL222. Differences between sub-populations gradually disappear over time but are still noticeable up to days after the last application of light to the population (Figure 2c). This is quite remarkable as it is difficult to comprehend, at a first glance, how a constitutively expressed gene can display a cellular memory of a stimulus that is retained over several tens of cell generations. In conclusion, cell-to-cell variability in EL222, which previously seemed to be negligible for the characterization of the system, suddenly appears to be of key importance for understanding how population dynamics emerge from the differentiation system. We may thus ask ourselves if a dedicated characterization of the system with a multi-scale stochastic kinetic model that takes into account both single-cell and population processes would have allowed us to retain predictable functionality.

### Single cell modeling of the integrated differentiation system

To test if consequences of changes in the system can be understood and predicted, we constructed a multi-scale stochastic kinetic model of the integrated differentiation system and a model of plasmid copy number fluctuations and asked if the models can be composed to predict emerging single-cell and population dynamics when the differentiation system is expressed from plasmids. Concretely, since variability in EL222 appeared to be of key importance, we deployed a model of bursty production of EL222 and cell differentiation to represent the differentiation system:

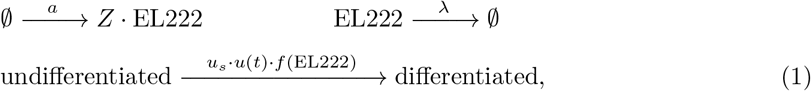

where *u_s_* is the maximal single cell differentiation rate for given fixed light intensity, *u*(*t*) is equal to one in the presence of light and equal to zero otherwise, *λ* is the cells’ growth rate, and *a* is the rate at which protein bursts occur. Protein production bursts are of size *Z* and assumed to be geometrically distributed with average burst size *b*, 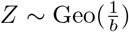, as dictated by classical results for modeling stochastic gene expression [18, 19]. To keep the model as simple as possible, we neglected possible delays or noise caused by the production and action of recombinase or the experimental detection of recombined cells. Instead, we assumed that the probability per unit time for a cell to differentiate in the presence of light is directly a function of the amount of EL222, *f* (EL222), in the cell.

When *u*(*t*) = 0, the EL222 distribution converges approximately to a stationary Gamma distribution that is determined by average burst size and frequency [18, 19]. Growing cells in the dark and measuring their mVenus fluorescence by flow cytometry then allows one to determine burst size and frequency (up to a fluorescence scaling factor) from mean and coefficient of variation of measured fluorescence distributions (Figure 3a, left). When light is applied to the population, *u*(*t*) = 1, the dynamics of the differentiated population fraction are determined by the specific shape and parameters of the differentiation function *f* (EL222). Matching emerging population dynamics of the integrated system for the light pattern in Figure 1d, we find that a steep Hill-function with a threshold significantly larger than average amounts of EL222 leads to good agreement with the data (Figure 3a, right and Supporting Information, Section S.2 A).

**Figure 3:**
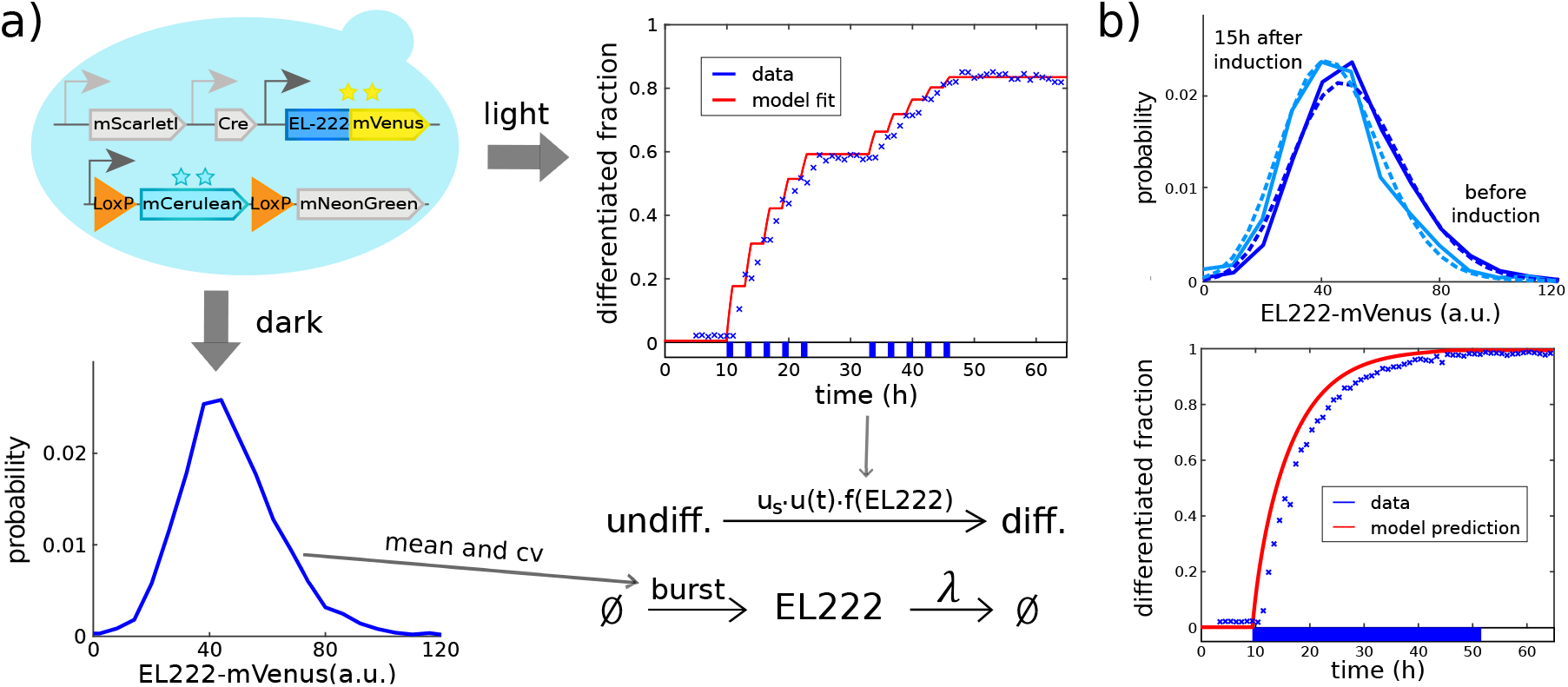
(a) Characterization of the differentiation system. *Bottom:* Extracting mean (= 50.6a.u.) and coefficient of variation (= 0.35) from measured stationary EL222 distributions before light induction allows one to determine burst frequency and average burst size of the EL222 production model up to a fluorescence scaling factor (Supporting Information, Section S.2). *Right:* Exposing cell populations to light pulses and measuring dynamics of the resulting population composition (crosses: data, line: model fit, bottom: applied light sequence) then allows one to determine parameters of the differentiation function *u_s_* · *f* (EL222) (Supporting Information, Section S.2). **(b) Dynamics in continuous light**. *Bottom:* Exposing cells to continuous light eventually leads to differentiation of the entire population. *Top:* Cells that remain undifferentiated after light induction (light blue) seem to be characterized by slightly lower EL222 levels compared to initial levels (dark blue) according to both model predictions (dashed) and data (solid). Data from further time points is provided in the Supporting Information, Figure S.2.

Mathematically, the model couples a variability generating process (stochastic production of EL222) to a population process that selectively differentiates cells based on their phenotypes. The coupling of single-cell and population scale tells us that, in response to light, we should expect preferential differentiation of cells with high levels of EL222. Correspondingly, differentiated cells should display increased amounts of EL222 shortly after light while the (sub-)population distribution of undifferentiated cells should shift to lower levels. However, this selective population split is counter-acted by the fact that the same variability generating process is operating in all cells and that this process will always drift to the original EL222 distribution in both sub-populations at a time scale that is determined by the cells’ growth rate (see Supporting Information, Section S.2 B for details). In the absence of light, we therefore expect distributions to revert back to the original EL222 distribution while the continuous presence of light should lead to a maximal shift of the distribution to a quasi-stationary condition in which single cell and population process are dynamically balanced until eventually all cells will have differentiated. The quasi-stationary EL222 distribution for undifferentiated cells in the presence of light can be obtained via an eigenvector calculation from a non-linear master equation that can be derived from the original master equation of the bursty protein production model (see Supporting Information, Section S.2 B). Performing this calculation, we find that the maximum possible shift in EL222 distributions of undifferentiated cells that can potentially be observed in experiments is fairly small and of similar size as experimental errors due to inaccurate deconvolution or reactor-to-reactor variability. To test whether such a shift can nevertheless be detected, we exposed cells to continuous light and collected measurements at time points early enough after induction such that sufficiently many cells remain undifferentiated to allow for reliable quantification of EL222 distributions. We find that experimental EL222 distributions of undifferentiated cells indeed seem to show a small shift towards lower levels in response to continuous light (Supporting Information, Figure S.2). This shift is in good agreement with distribution dynamics predicted from the model (Figure 3b, top and Supporting Information, Figure S.4). As a side note, the model provides a very good prediction of the increase in the differentiated fraction in response to continuous light (Figure 3b, bottom). Despite the possible presence of small selection effects, we can overall conclude that sufficiently low noise in EL222 production coupled to sufficiently fast fluctuations implies that cell-to-cell variability has only small consequences for emerging population dynamics. It is now clear, however, that this conclusion will change if either noise levels or time scales of the single cell process are modified.

### Consequences of plasmid copy number fluctuations

In addition to a single-cell model of the differentiation system, we require a model that captures cell-to-cell variability in plasmid copy numbers. Many, often detailed, models of plasmid copy number fluctuations exist in the literature [20, 21]. In order to keep the system characterization as simple as possible, we decided to omit any detailed mechanistic description of processes such as replication failure or unequal division of plasmids between mother and daughter cell [22]. Instead, we chose to represent plasmid copy number fluctuations by a simple stochastic birth-death process with both birth rate (representing replication) and death rate (representing dilution due to cell growth) being linear in the plasmid copy number:

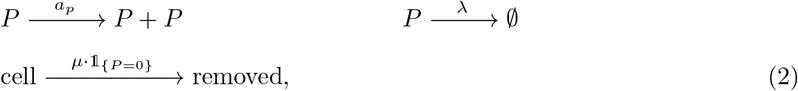

where *a_p_* is the plasmid replication rate, *λ* is the growth rate of cells, and *μ* is the rate at which cells that have lost the plasmid 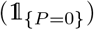 are removed from the population when cells are growing in selective media. Replication failure can be implicitly incorporated by choosing the replication rate smaller than the cells’ growth rate, which is in any case a necessary feature of a birth-death process model since expected plasmid copy numbers in cells would diverge to infinity if plasmids are replicated faster than cells divide. For *a_p_* < *λ* (and *μ* = 0), on the other hand, the process will eventually reach zero plasmids with probability one. The rate at which cells in the population lose the (last copy of the) plasmid is then determined by the difference between replication and growth rate. When selective media is used for growth, *μ* > 0, cells that have lost the plasmid will either die or be outgrown and the plasmid copy number distribution of the population will remain stable. However, this neither implies that there exist no cells without plasmids in the population nor do plasmid copy numbers remain constant from the perspective of single cells. Instead, we should expect a dynamic equilibrium and a quasi stationary plasmid copy number distribution in which the single cell plasmid loss process is balanced with selective removal of cells without plasmids (Supporting Information, Section S.3 B). From a mathematical perspective, this result is equivalent to what was obtained previously for EL222 distributions in the undifferentiated cell population. In both cases, a variability generating process at the single-cell scale (EL222 fluctuations vs. plasmid copy number fluctuations) is coupled to a state-dependent removal process at the population scale (differentiation vs. removal of cells that have lost the plasmid) and leads to the same type of non-linear master equation for the cells that have not been removed yet (compare Supporting Information, Sections S.2 B and S.3 B).

To characterize the full multi-scale dynamics of plasmid copy numbers and cell populations, we switched cells from selective to non-selective media and measured how the average abundance of a constitutively expressed protein decays over time. We found that average fluorescence decays approximately exponentially with a rate that is 15% of the cells’ growth rate *λ* (Figure 4a, bottom). Mathematical analysis of the single-cell model (Supporting Information, Section S.3 B) shows that this is a direct consequence of the plasmid replication rate, *a_p_*, being 15% smaller than the cells’ growth rate *a_p_* = 0.85*λ*. Somewhat counter-intuitively, the net population growth rate in selective media is independent of *μ*. Faster removal of cells that have lost the plasmid means faster reduction (or slower increase) in the total number of cells but this is compensated by the fact that, in stationary growth conditions, the population fraction of cells without plasmids becomes smaller when their removal rate is increased. Therefore, to determine *μ*, stationary growth rate measurements are insufficient. Instead, it is necessary to directly measure what fraction of the population has plasmids in stationary growth conditions since it can be shown (Supporting Information, Section S.3 A) that this fraction must be equal to 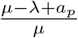 for *a_p_* < *λ* (plasmid numbers do not diverge) and *μ* > *λ* – *a_p_* (cells with plasmids can be maintained in the population). Correspondingly, we performed a colony counting experiment (see Supporting Information, Section S.3 A) and observed that approximately one third of all cells do not contain plasmids in stationary growth conditions (Figure 4a, top), which allowed us to determine that the removal rate of cells without plasmids, *μ*, must be approximately 45% of the cells growth rate, *μ* = 0.45*λ*. Together, the parameters *a_p_*, *μ* and *λ* completely characterize the multi-scale dynamics of plasmid copy number fluctuations and growth in selective media.

**Figure 4:**
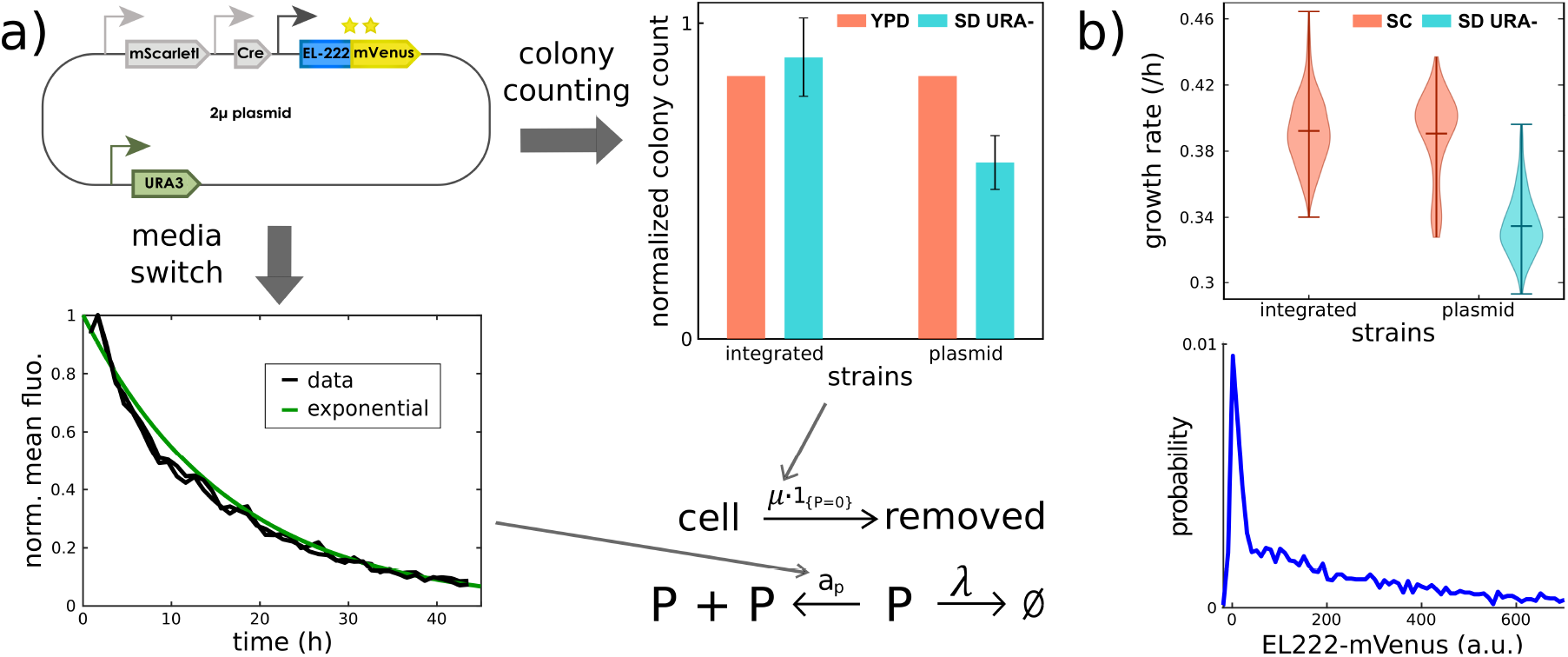
(a) Characterization of plasmid copy number dynamics. *Bottom:* After switching cells from selective to non-selective media, the rate of exponential decay of average levels of a protein that is constitutively expressed from plasmids (here EL222-mVenus) can be measured to determine the population plasmid loss rate and to fix the plasmid replication rate in the model. Mean fluorescence (normalized by its maximum) of two replicates is shown. *Right:* Stationary fractions of cells without plasmids determine the removal rate in selective media of cells that have lost the plasmid. These fractions can be measured by a colony counting experiment as described in the Supporting Information, Section S.3 A. **(b) Consequences of plasmid copy number fluctuations**. Expressing the differentiation system from plasmids leads to modified population distributions of EL222 (bottom) as well as changes in the effective population growth rate (top). Violin plots are composed of growth rate estimates based on cell density averaged over all reactors of the turbidostat and multiple experiments (in total 19 reactors for the strain with the integrated system, 12 for the strain with the plasmid-based system in selective media and 6 in non-selective media).

To test the model and to better understand possible consequence of plasmid copy number fluctuations, we experimentally determined effective population growth rates of our cells in selective and non-selective media. We found that, after pre-culture in selective media, populations of cells carrying the plasmid-based differentiation system quickly adopt the same growth rate as populations of cells carrying the integrated system version when transferred to non-selective media. Subsequently, the population growth rate remained constant and no dependence on the fraction of cells that still carry plasmids was detected. When grown in selective media, however, populations of cells carrying the plasmid-based differentiation system display an effective growth rate that is reduced by approximately 15% (Figure 4b, right). Analyzing the model, we find that the reduction in growth is a consequence of removal of cells that have lost the (last copy of) the plasmid and that the observed reduction by 15% is in quantitative agreement with model predictions and another consequence of the plasmid replication rate, *a_p_*, being 15% smaller than the single-cell growth rate *λ*. Overall, we should expect that placing the differentiation system on plasmids will significantly alter variability in EL222 levels (Figure 4b, bottom) and also modify effective population growth.

Having constructed single cell models of the differentiation system and plasmid copy number fluctuations, we are now in a position to ask the central question of this manuscript: is it possible to quantitatively predict functionality of the plasmid-based differentiation system without having to change the model or even just to re-identify model parameters for the new conditions?

### Parameter-free prediction of circuit functionality

To test if the dynamics of the plasmid-based differentiation system can be predicted by combining models calibrated on parts of the system, we coupled the single-cell model of the integrated differentiation system, Eq.(1), to the model of plasmid copy number fluctuations, Eq.(2), to obtain a composed model that can be stated in reaction network form as

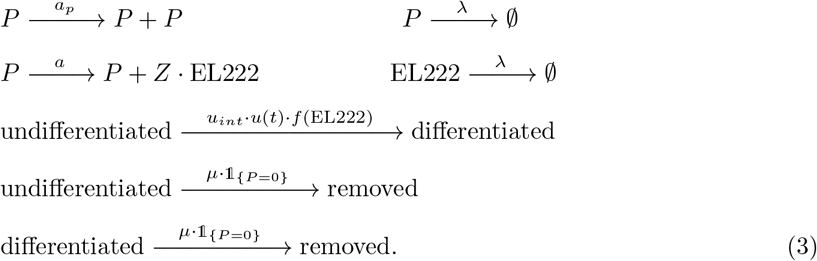

We highlight that the coupling of the two models is very natural and does not introduce any new parameters: the protein burst rate is now scaled by the plasmid copy number since production can occur from any copy of the plasmid while all other model parts are exactly the same as previously characterized. To test if complex functionalities of our circuit such as population dynamics emerging from single-cell stochastic biochemical processes can be predicted without any adjustment of model parameters, we used the composed model to predict the dynamics of the plasmid-based version of the differentiation system for the light stimulation pattern in Figure 2d and compared the results to data. We find that population dynamics of differentiated and undifferentiated cells are very well predicted without any adjustment of model parameters (Figure 5b).

**Figure 5:**
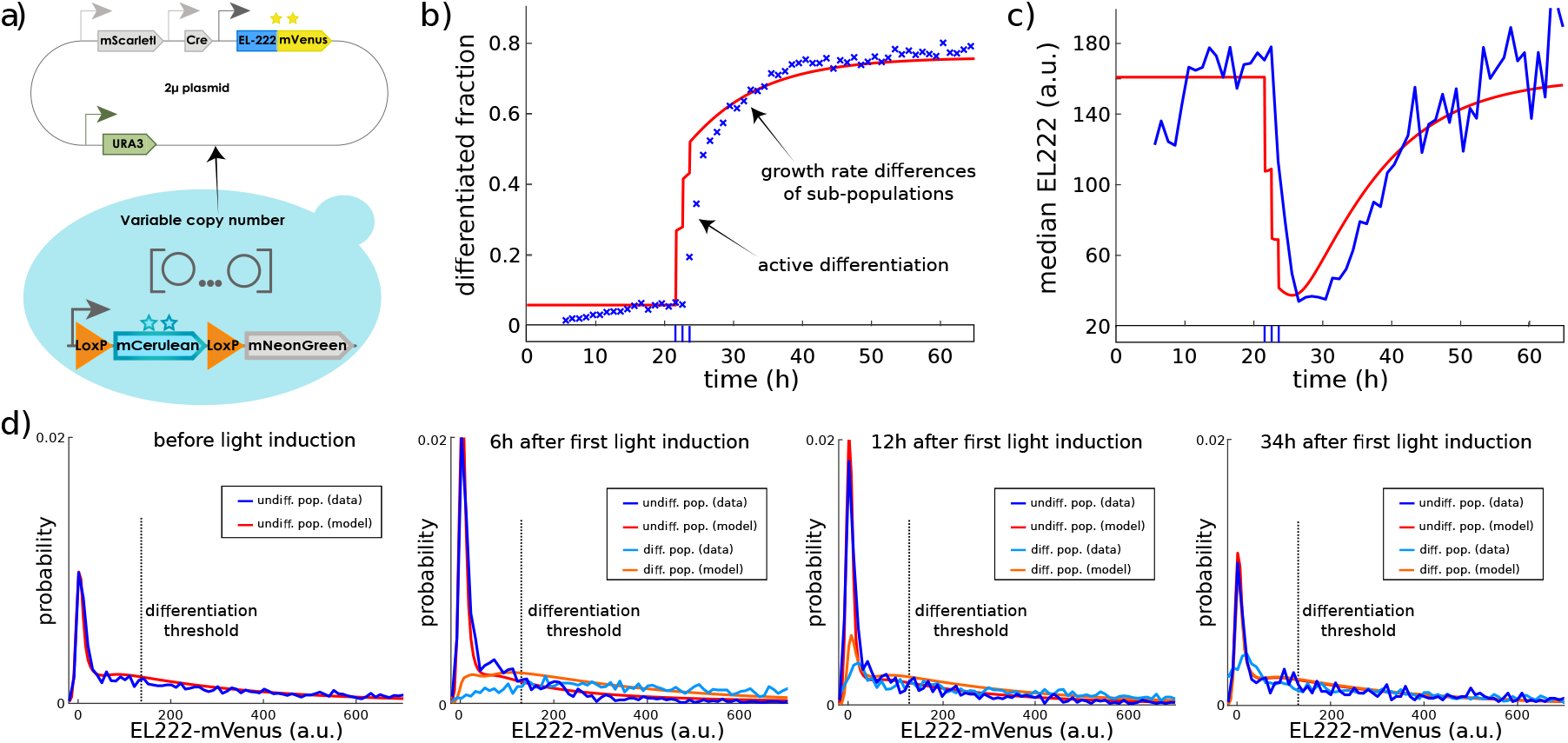
Parameter-free prediction of emerging population dynamics for the plasmid-based differentiation system. **(a)** Sketch of the plasmid-based differentiation system. **(b)** Emerging dynamics of the differentiated fraction according to the composed model (red) for the same light input as in Figure 2d are compared to experimental data (blue). According to the model, late increases in the differentiated fraction are due to varying sub-population growth rates and not caused by active differentiation. **(c)** Upon light induction, median EL222 levels of undifferentiated cells drop 5-6 fold as predicted by the composed model. **(d)** *Left panel:* The model-predicted EL222 distribution in the dark agrees very well with experimental data even though this data was not used to parametrize the model. According to the model, the peak of the distribution close to zero corresponds to the 33% of cells without plasmids (see Figure 4a). *Middle panels:* Shortly after the application of light, cells with EL222 levels above the threshold parameter of the differentiation Hill function (dashed line) are enriched in the differentiated population while the undifferentiated population contains more cells without plasmids (increased peak close to zero). *Right panel:* When cells are subsequently kept in the dark, EL222 distributions converge back to the initial condition. Model predictions show remarkable agreement with measured EL222 distribution dynamics, except for seemingly lower numbers of differentiated cells without plasmids in the data (in particular after 34h). This small mismatch is presumably caused by inaccuracies in deconvolution (the presence of mNeonGreen in cells makes it difficult to precisely quantify low mVenus levels in cells, Supporting Information, Section S.7). Full time-varying distribution data of all fluorescent reporters for this experiment is displayed in the Supporting Information, Figure S.8.

Comparing EL222 distributions in the two sub-populations between model and data shows that the high quality of population-level predictions is a consequence of the fact that the model predicts correctly how single-cell processes will operate in union with population-level processes to shape the full dynamics of EL222 distributions in undifferentiated cells (Figure 5c,d and Supporting Information, Figure S.5). In particular, the heavy tails of EL222 distributions for cells growing in darkness (observed already in Figure 2b) emerge naturally from the fact that plasmid copy numbers can fluctuate significantly in the model. These heavy tails imply that significantly more cells have EL222 levels above the threshold of the differentiation Hill function *f* (EL222) and will recombine quickly upon light induction. Shortly after light induction, the remaining undifferentiated cell population is therefore shifted to significantly lower EL222 levels (5-6 fold reduction of the median, Figure 5c) while the differentiated cell population inherits the heavy tail of the original distribution (Figure 5d). While not experimentally measurable, it can be deduced that according to the model the same holds true for plasmid copy number distributions in sub-populations (Supporting Information, Figure S.6). As a consequence, the population differentiation rate spikes very high upon first light induction but is significantly reduced when subsequent light pulses are applied before the EL222 distribution of the undifferentiated cell population has converged back to its initial condition (Supporting Information, Figure S.7). When the light stimulus is maintained for some time, fluctuations in plasmid copy numbers create larger fluctuations in EL222 amounts compared to the integrated version of the system, which leads to more frequent threshold crossing events and therefore larger population differentiation rates.

Eventually, the plasmid-based version of the system reaches differentiated population fractions close to 100% very quickly when light is maintained (Supporting Information, Figure S.10) despite the fact that a large part of the population (around a third) displays mVenus levels that are close to zero at all time points. According to the model, these are cells that have lost the plasmid (Supporting Information, Figure S.6), cannot differentiate, but are likely to be removed at subsequent time points due to growth in selective media. The coupling of plasmid loss dynamics and selective media with the differentiation system therefore leads to unintuitive population dynamics in which seemingly the entire population can recombine despite a continuous presence of many cells that are not carrying any copy of the differentiation system. Another complex consequence of the coupling of single-cell and population processes is that the split in plasmid copy numbers between sub-populations that follows from selective differentiation of cells with high EL222 levels leads to different sub-population growth rates in selective media (Supporting Information, Figure S.7) since cells that have lost the plasmid (or are close to losing it) are enriched in the undifferentiated subpopulation. This implies that the differentiated population fraction will continue to increase even if light is removed and no more active differentiation takes place, which explains why the assumption of the simple deterministic model in Figure 1d, that the differentiated population fraction can only increase due to active differentiation, led to structural incapacity of the model to explain the slow transient differentiation dynamics and significantly increasing differentiated fractions up to 10h after last light induction (Figure 5b). If the light stimulus is removed, EL222 sub-population distributions converge back to the original distribution (Figure 5d) but remain noticeably different for at least a day after last light induction. This experimental observation is in good agreement with slow convergence of sub-population plasmid copy numbers to their quasi-stationary distribution in selective media, which creates a surprisingly long sub-population memory of applied light stimuli (Supporting Information, Figure S.6). We conclude that deploying multi-scale stochastic chemical kinetics models for understanding the interplay of single-cell and population processes allows us to understand and predict complex emerging dynamics when the differentiation system is placed on plasmids and cells are grown in selective media. Population dynamics for plasmid strains, despite being shaped by cell-to-cell variability, are deterministically reproducible (Supporting Information, Figure S.12). Thus, the capacity to predict such dynamics implies that the interplay of single-cell and population processes can be exploited for creating features of microbial community dynamics that would otherwise be difficult to engineer.

### Optogenetic control of constitutive gene expression

In the previous section, we demonstrated that coupling of single-cell and population processes may lead to outcomes that are fairly unintuitive such as dynamically changing population distributions of a constitutively expressed gene or increasing fractions of differentiated cells in the absence of active differentiation. In practice, such couplings will often be seen as a nuisance but we may also ask if the interplay of stochastic single-cell processes and population dynamics can be exploited to create and control features of cell populations that would otherwise be impossible to engineer. For instance, in light of the results of this paper, we may ask if it is possible to use light to regulate constitutive gene expression, plasmid copy numbers, and growth rates of sub-populations via targeted differentiation of cells with high EL222 levels. Since neither plasmid copy numbers nor sub-population growth rates are directly measurable on our experimental platform, we set ourselves the goal to regulate the constitutively expressed EL222. Concretely, according to our theoretical results (convergence to a quasi-stationary distribution, Supporting Information, Sections S.2 B and S.3 B), it should be possible to maintain EL222 levels in the undifferentiated cell population at reduced constant levels by applying continuous light for sufficiently long. Indeed, we find that EL222 distributions in undifferentiated cells quickly reach a distribution that remains invariant when the population is exposed to continuous light (Figure 6c, left). However, this distribution is characterized by a median of almost zero (Figure 6b), which indicates that only cells that have no EL222 (and presumably no plasmids) remain undifferentiated. Furthermore, 15 − 20h after the start of light application, almost the entire population is differentiated. To test if also low but non-zero EL222 levels in undifferentiated cells can be stably maintained, we exposed cells to a number of different light sequences that mimic continuous light with short pulses that are regularly repeated with an interpulse duration that is short in comparison to the duration of the experiment. We find that applying light for 2min every 4h or for 1min every 1h leads to approximately halved median EL222 levels in undifferentiated cells (Figure 6b). In both cases, the full EL222 distribution of undifferentiated cells remains approximately invariant after around 12h have passed from first light application (Figure 6c, middle, see also Supporting Information, Figure S.13). Finally, we applied a light sequence with only half-minute light pulses applied every 4h and found that, in this case, the EL222 distribution of undifferentiated cells differs only very moderately from the initial population distribution at all time points (Figure 6c, right) despite the fact that the differentiated population fraction increases (Figure 6a).

**Figure 6:**
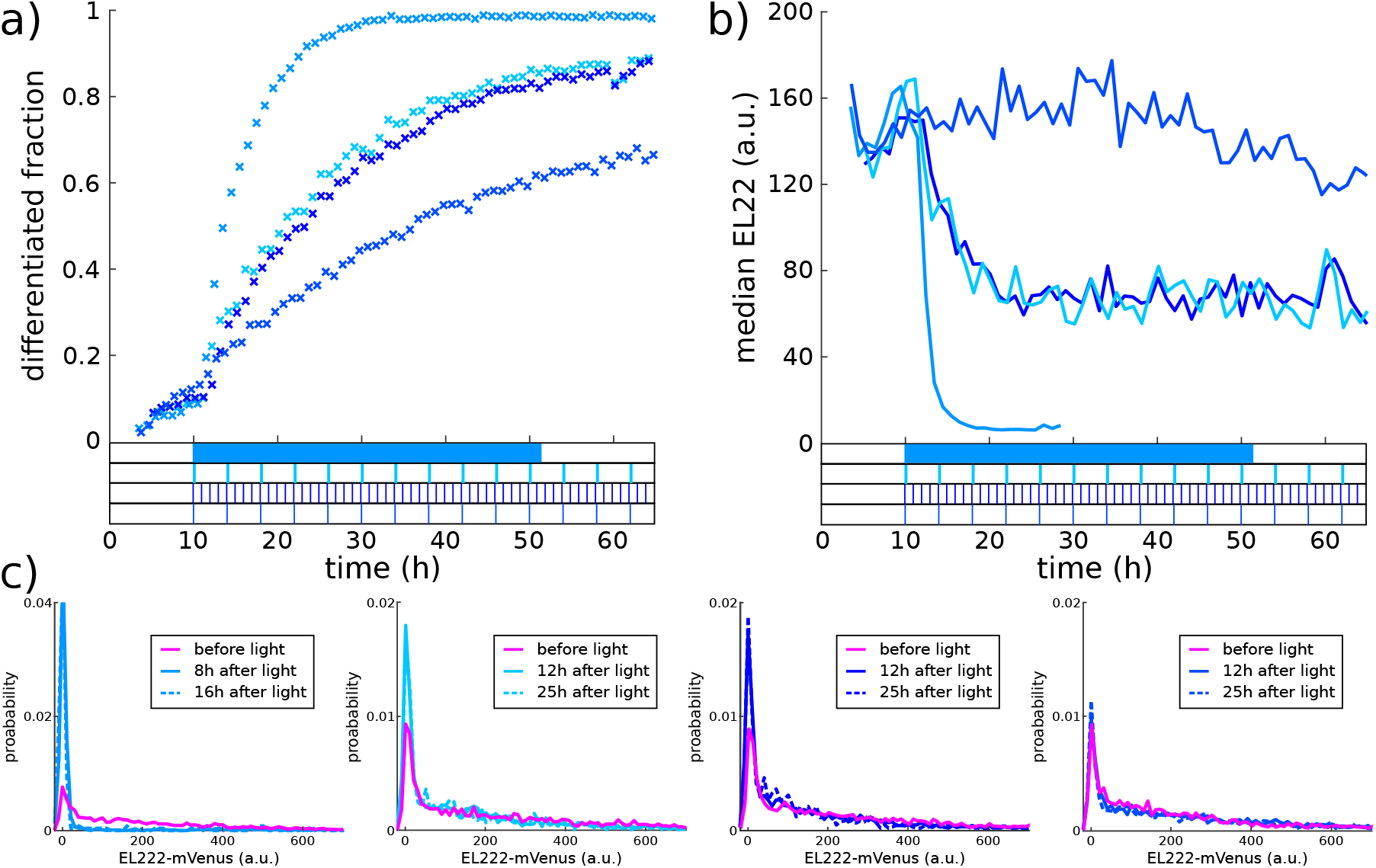
Optogenetic control of constitutive gene expression. **(a)** Regularly repeated short light pulses (shown at the bottom) lead to a slow steady increase in the differentiated population fraction while the differentiated fraction increases fairly quickly in response to continuous light. **(b)** Over intermediate time periods, repeated selective differentiation and reversion to the mean balance each other, which leads to approximately constant EL222 medians in all cases. **(c)** After transient dynamics in response to the start of the light sequence, EL222 distributions in undifferentiated cells reach a quasi-stationary form that remains invariant for extended time periods (see also Supporting Information, Figure S.13). Color coding of distributions labels correspondence to the light sequences and data in panels a and b.

To conclude, understanding and characterizing the consequences of cell-to-cell variability allowed us to use light to optogenetically regulate the (sub-population) level of a constitutively expressed gene, a feat that seems quite counter-intuitive at a first glance and that is not realizable in any obvious way by other means.

## Discussion

Quantitatively predicting the dynamics of complex synthetic circuits before the circuit is constructed is the key challenge that needs to be mastered to turn synthetic biology into a true engineering discipline. Yet, while our capacity to construct complex circuits is continuously increasing, our capability to predict circuit functionality from supposedly known and characterized circuit components remains, at best, limited to very tightly constrained operating conditions and qualitative and/or stationary outputs [23]. We exemplified this problem for the case of our optogenetic differentiation system by showing that predictions obtained from a simple deterministic model break down as soon as the context in which the system is used (plasmid-based expression of proteins and growth in selective media) is modified (Figure 2). The concrete reasons for the failure of model predictions may be manifold and caused by unexpected component-to-component interactions (e.g. retroactivity [24] or resource competition [25]) or couplings of the circuit to processes of the host (e.g. burden [11] or saturation of the host’s degradation machinery [4]). Eventually, however, all these reasons are but different facets of a common problem: our incapacity to foresee the consequences of complex inter-dependencies of a circuit in vivo.

For our optogenetic differentiation system, expressing proteins from plasmids led to unforeseen couplings of plasmid copy number fluctuations, growth in selective media, and selective differentiation, which, for instance, created seemingly slow “differentiation dynamics” that remained mysterious (even to us) in the absence of an explanatory model (Figure 2d). We thus set out to construct dedicated multi-scale stochastic kinetic models of the circuit’s components in their population context (Figures 3 and 4). Since the chemical master equation (CME) in its standard form does not capture population processes, it was necessary to augment the CME and to derive a non-linear version for conditional probabilities to correctly capture dynamics of population distributions (Supporting Information, Sections S.2 B and S.3 B). We then used our theoretical results to determine experiments that can be used to readily extract parameters of the two component models from data (Figures 3a and 4a). Merging the so obtained models to construct a model of the plasmid-based differentiation system (Eq.[3]), we found that the composed model does not only resolve the previously not understood features but quantitatively predicts the consequences of complex component interactions without any adjustment of model parameters (Figure 5). This is a particularly encouraging finding as it demonstrates that, at least for the system studied in this paper, our incapacity to foresee the consequences of complex inter-dependencies of circuit components and couplings of single-cell and population processes can be remedied by appropriate characterization of circuit components. Importantly, with the ability to foresee consequences comes the possibility to exploit such couplings in the design [26]. This is evidenced by our result that the circuit allows one to modulate constitutive gene expression (and presumably plasmid copy numbers and growth rates) in sub-populations via the application of light (Figure 6) - a feat that seems impossible upon a first glance at the circuit’s wiring diagram (Figure 1a).

To conclude, we highlight that for the application considered in this paper, the crucial ingredient for obtaining a faithful model was to augment the CME to incorporate population processes in addition to single-cell dynamics that are classically tracked in stochastic models. It is to be expected that the same will hold true for many other applications where couplings of single-cell and population processes are likely to be at play [27, 28, 29, 30, 31]. This is notably the case for synthetic circuits that are constructed to produce proteins in large quantities since this creates a burden for cells that may lead to growth rate variability between cells and a dynamical enrichment of low-producing cells upon induction of protein production. Furthermore, similar couplings between scales are likely also present in many natural systems such as selective killing of cancer cells with particular states of internal processes in response to treatments that induce the apoptotic pathway [32] or differential responses of bacteria to antibiotic treatments [33] to name but two possible examples. We therefore expect that our approach for calculating with multi-scale stochastic kinetic models will be of use far beyond the particular case study considered here. It needs to be noted, however, that for cases where the single-cell model is more high-dimensional than the fairly small models considered here, tracking the entire solution of the corresponding master equation will be computationally infeasible. Further work is thus necessary to develop and test approaches for approximately calculating with multi-scale stochastic kinetic models [34, 14].

## Author contributions

C.A. performed experimental work. J.R. performed modeling and mathematical work. C.A. and J.R. analyzed data. F.B. helped with experimental platforms and data analysis. G.B. and J.R. conceived and supervised the project. All authors wrote the paper.

## Acknowledgements

We thank Sebastian Sosa Carrillo for providing plasmid backbones and Zachary Fox for providing software for efficient master equation solving. C.A. is enrolled in the Frontières de l’Innovation en Recherche et Education (FIRE) doctoral school hosted by Université de Paris. This work was supported by the H2020 Fet-Open COSYBIO grant (grant agreement no. 766840) and by the ANR grant CyberCircuits (ANR-18-CE91-0002).

## Data, material and code availability

Data can be found on Zenodo (DOI:10.5281/zenodo.5155290), and code used for data analysis in the GitLab repository (https://gitlab.inria.fr/InBio/Public/predicting-selection-effects). Plasmids and strains are available from the authors upon request.

## References

[1] Elowitz M, Levine A, Siggia E, Swain P (2002) Stochastic gene expression in a single cell. Science 297:1183–1186.

[2] Kwok R (2010) Five hard truths for synthetic biology. Nature 463:288–290.

[3] Del Vecchio D, Qian Y, Murray R, Sontag E (2018) Future systems and control research in synthetic biology. Annual Reviews in Control 45:5–17.

[4] Potvin-Trottier L, Lord N, Vinnicombe G, Paulsson J (2016) Synchronous long-term oscillations in a synthetic gene circuit. Nature 538:514.

[5] Lugagne JB, et al. (2017) Balancing a genetic toggle switch by real-time feedback control and periodic forcing. Nature Communications 8:1671.

[6] Raser J, O’Shea E (2005) Noise in gene expression: Origins, consequences, and control. Science 309:2010–2013.

[7] Zechner C, et al. (2012) Moment-based inference predicts bimodality in transient gene expression. Proceedings of the National Academy of Sciences of the USA 109:8340–8345.

[8] Munsky B, Neuert G, van Oudenaarden A (2012) Using gene expression noise to understand gene regulation. Science 336:183–187.

[9] Neuert G, et al. (2013) Systematic identification of signal-activated stochastic gene regulation. Science 339:584–587.

[10] Milias-Argeitis A, Rullan M, Aoki S, Buchmann P, Khammash M (2016) Automated optogenetic feedback control for precise and robust regulation of gene expression and cell growth. Nature Communications 7:12546.

[11] Weiße A, Oyarzún D, Danos V, Swain P (2015) Mechanistic links between cellular trade-offs, gene expression, and growth. Proceedings of the National Academy of Sciences of the USA 112:E1038–E1047.

[12] Gillespie D (1992) A rigorous derivation of the chemical master equation. Physica A 188:404–425.

[13] Schnoerr D, Sanguinetti G, Grima R (2017) Approximation and inference methods for stochastic biochemical kinetics - a tutorial review. Journal of Physics A: Mathematical and Theoretical 50:093001.

[14] Lunz D, Batt G, Ruess J, Bonnans J (2021) Beyond the chemical master equation: stochastic chemical kinetics coupled with auxiliary processes. PLoS Computational Biology 17:e1009214.

[15] Aditya C, Bertaux F, Batt G, Ruess J (2021) A light tunable differentiation system for the creation and control of consortia in yeast. bioRxiv.

[16] Baumschlager A, Aoki S, Khammash M (2017) Dynamic blue light-inducible t7 rna polymerases (opto-t7rnaps) for precise spatiotemporal gene expression control. ACS synthetic biology 6:2157–2167.

[17] Bertaux F, et al. (2020) Enhancing bioreactor arrays for automated measurements and reactive control with reacsight. bioRxiv.

[18] Friedman N, Cai L, Xie S (2006) Linking stochastic dynamics to population distribution: An analytical framework of gene expression. Physical Review Letters 97:168302.

[19] Shahrezaei V, Swain P (2008) Analytical distributions for stochastic gene expression. Proceedings of the National Academy of Sciences of the USA 105:17256–17261.

[20] Paulsson J, Ehrenberg M (2001) Noise in a minimal regulatory network: plasmid copy number control. Quarterly Reviews of Biophysics 34:1–59.

[21] Gnügge R, Liphardt T, Rudolf F (2016) A shuttle vector series for precise genetic engineering of saccharomyces cerevisiae. Yeast 33:83–98.

[22] Huh D, Paulsson J (2011) Non-genetic heterogeneity from stochastic partitioning at cell division. Nature Genetics 43:95–100.

[23] Nielsen A, et al. (2016) Genetic circuit design automation. Science 352:aac7341.

[24] Del Vecchio D, Ninfa A, Sontag E (2008) Modular cell biology: retroactivity and insulation. Molecular systems biology 4:161.

[25] Qian Y, Huang H, Jimnez J, Del Vecchio D (2017) Resource competition shapes the response of genetic circuits. ACS Synthetic Biology 6:12631272.

[26] Hasty J, Pradines J, Dolnik M, Collins J (2000) Noise-based switches and amplifiers for gene expression. Proceedings of the National Academy of Sciences of the USA 97:2075–2080.

[27] Tan C, Marguet P, Lingchong Y (2009) Emergent bistability by a growth-modulating positive feedback circuit. Nature Chemical Biology 5:842–848.

[28] Shahrezaei V, Marguerat V (2015) Connecting growth with gene expression: of noise and numbers. Current Opinion in Microbiology 25:127–135.

[29] Ghusinga K, Dennehy J, Singh A (2017) First-passage time approach to controlling noise in the timing of intracellular events. Proceedings of the National Academy of Sciences of the USA 114:693–698.

[30] Ruess J, Pleška M, Guet C, Tkačik G (2019) Molecular noise of innate immunity shapes bacteria-phage ecologies. PLoS Computational Biology 15:e1007168.

[31] Miano A, Liao M, Hasty J (2020) Inducible cell-to-cell signaling for tunable dynamics in microbial communities. Nature Communications 11:1193.

[32] Bertaux F, Stoma S, Drasdo D, Batt G (2014) Modeling dynamics of cell-to-cell variability in trail-induced apoptosis explains fractional killing and predicts reversible resistance. PLoS Computational Biology 10:e1003893.

[33] Wakamoto Y, et al. (2013) Dynamic persistence of antibiotic-stressed mycobacteria. Science 339:91–95.

[34] Duso L, Zechner C (2020) Stochastic reaction networks in dynamic compartment populations. Proceedings of the National Academy of Sciences of the USA 117:22674–22683.

