## Supporting Information for "Using single-cell models to predict the functionality of synthetic circuits at the population scale"

#### S.1 A simple deterministic population model of differentiation.

Figure 1c,d in the main paper displays data collected in an experiment with the strain carrying the integrated version of our differentiation system. The data in panel c of that figure can also be displayed in the form of time-varying distributions of the four fluorescent reporter proteins (Figure S.1).

As stated in the main text, a simple deterministic model can be used to describe emerging population dynamics for the integrated version of the system fairly well. This model is given by the following ordinary differential equations

$$\begin{aligned} \frac{d}{dt}n_u &= \lambda n_u - u_{int}u(t)n_u \\ \frac{d}{dt}n_d &= \lambda n_d + u_{int}u(t)n_u, \end{aligned} \tag{1}$$

where $n_u$ is the (expected) number of undifferentiated cells, $n_d$ is the number of differentiated cells, $\lambda$ is the (single cell) growth rate and measurable as $\lambda = \frac{\ln(2)}{103}\mathrm{min}^{-1}$ for all our strains, $u(t)$ is equal to one when light is applied and zero otherwise, and $u_{int}$ is the single free parameter of the model and represents the differentiation rate for the fixed light intensity that was used in the experiments. For the results in Figure 1d (solid line) in the main text, we fixed $u_{int} = 0.168\mathrm{h}^{-1}$. We note that the integral of the differentiation rate in this model increases linearly with the time of light application. When very short light inputs are used, however, we observe that the differentiated fraction increases less than expected from the value of $u_{int}$ given above. This is presumably because some cells do not produce sufficient recombinase when light is applied only very briefly. Better agreement of the model with data for short light signals could be obtained by including a recombinase species in the model that starts to accumulate in response to light and triggers a positive rate of differentiation only when it has sufficiently accumulated. For the sake of simplicity, and in order to not introduce additional parameters into the model, we have decided to nevertheless neglect recombinase and

*contributed equally
†

to limit ourselves to sufficiently long light signals. We note that the same arguments apply to subsequent stochastic models of the integrated differentiation system since recombinase has been omitted also for these models, which structurally limits the models to sub-linear dependencies of the integrated differentiation rate on the duration for which light is applied.

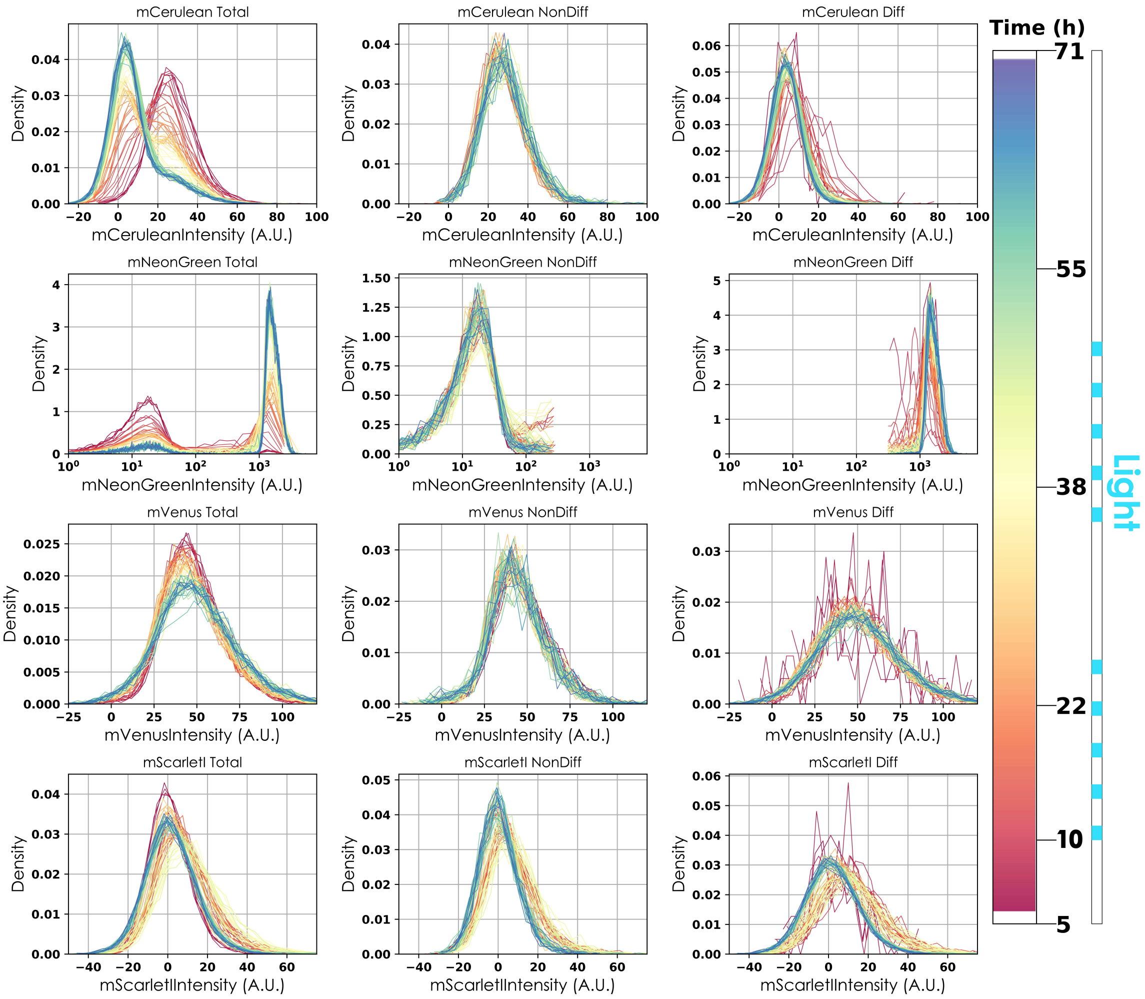


Figure S.1: **Distribution dynamics of the four fluorescent reporter proteins for the light sequence in Figure 1 in the main paper.** Color coding indicates the time of the experiment. Rows correspond to the different fluorescent reporters, columns to different sub-populations (left: all cells, middle: undifferentiated cells, right: differentiated cells) after classification based on mNeonGreen abundance. We note that due to changing sizes of sub-populations, fluorescence distributions need to be extracted from varying (sometimes low) cell numbers.

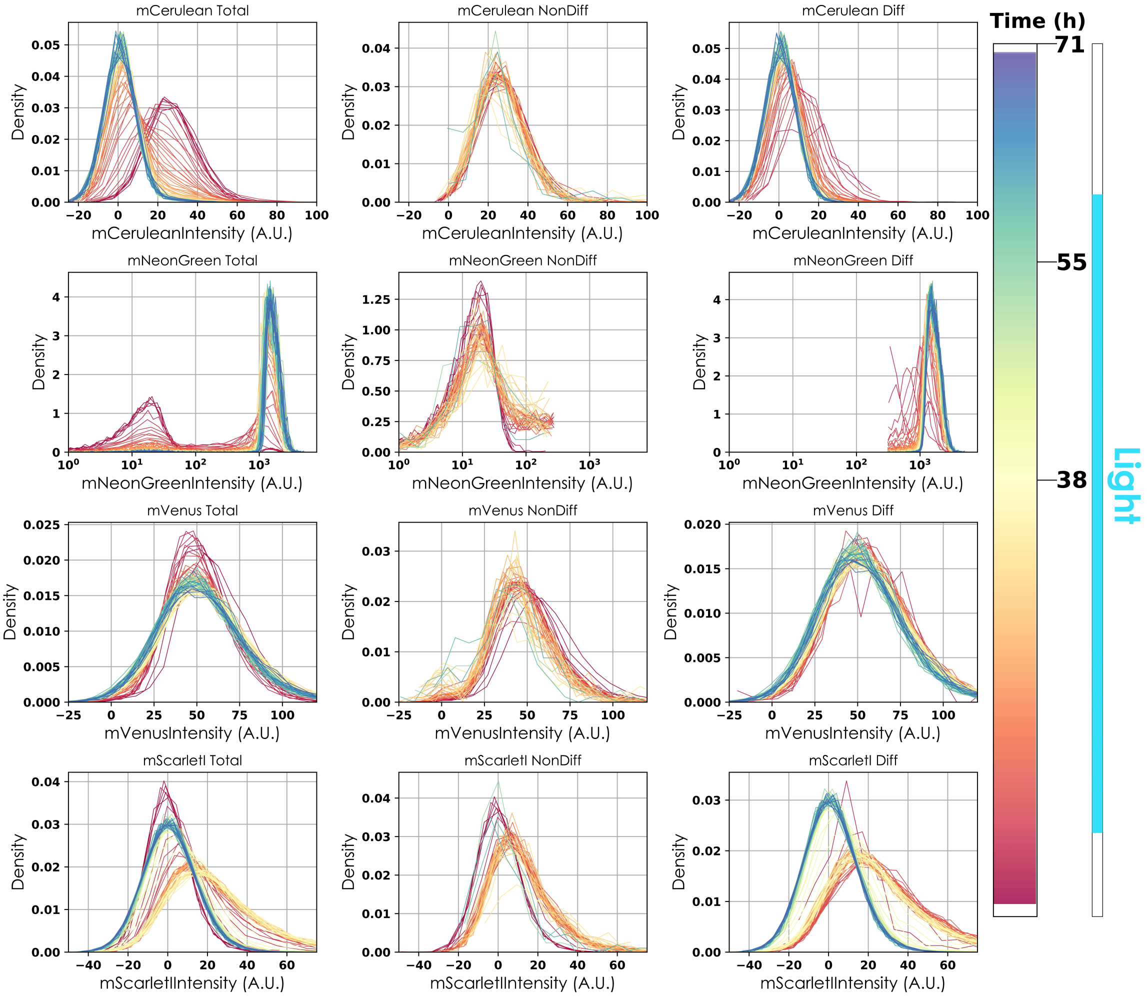


Figure S.2: **Distribution dynamics of the four fluorescent reporter proteins in response to continuous light.** Color coding indicates the time of the experiment. Rows correspond to the different fluorescent reporters, columns to different sub-populations (left: all cells, middle: undifferentiated cells, right: differentiated cells) after classification based on mNeonGreen abundance. We note that due to changing sizes of sub-populations, fluorescence distributions need to be extracted from varying (sometimes low) cell numbers.

To determine the importance of cell-to-cell variability, we exposed cells to continuous light and measured if EL222 distributions of undifferentiated cells change due to selection effects in response to light. We note that small selection effects can, in principle, also be detected by comparing undifferentiated and differentiated cells shortly after light induction. However, differentiation replaces mCerulean by mNeonGreen in cells and small differences between sub-population might also be caused by variations in the accuracy of deconvolution in the presence of different fluorophores. Comparisons of EL222 distributions before and after light within the same sub-population therefore provide a more reliable approach to detect small selection effects. The full time-varying distribution data of all fluorescent reporters is displayed in Figure S.2. The panel for mVenus levels (3rd row) in undifferentiated cells (2nd column) shows that differences in EL222 distributions before and after light are noticeably present but overall quite small.

To test if the model in Eq.(1) can be used to predict differentiation dynamics for the plasmid-based version of the system, we used mVenus fluorescence to quantify mean constitutive gene expression from plasmids and found that it is 5.84 times larger compared to expression levels measured in the integrated strain. Correspondingly, the plasmid-based version of the differentiation system leads to larger differentiated fractions for the same light stimulation (Figure S.3). Multiplying $u_{int} = 0.168\text{h}^{-1}$ by this factor gives the dynamics in Figure 2d, dashed line, in the main paper. Finally, the dashed-dotted dynamics in Figure 2d of the main paper has been obtain by hand-tuning $u_{int}$ such that final stationary differentiated fractions are well-matched. Concretely, we used the value $u_{int} = 5.4\text{h}^{-1}$.

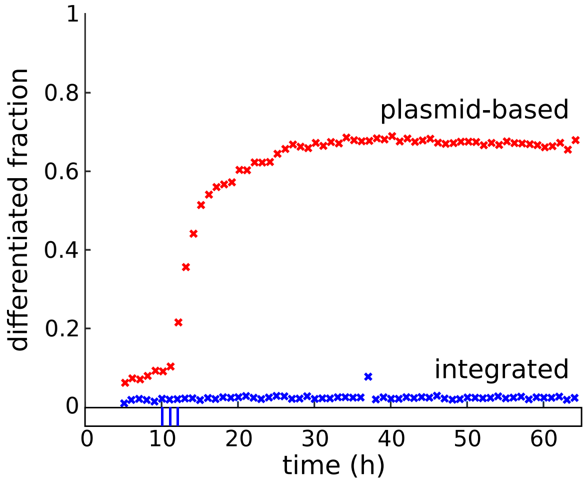


Figure S.3: Differentiation dynamics in response to the same light stimulation for plasmid-based and integrated differentiation system show large differences.

#### S.2 Stochastic modeling of the integrated differentiation system.

##### S.2.1 Construction of the model.

To characterize the functionality of the differentiation system at the single cell level, we deployed a one-dimensional model of bursty production of EL222 transcription factor coupled to a differentiation function that determines the probability per unit time for a cell to differentiate in the presence of light given its internal amount of transcription factor:

$$\emptyset \xrightarrow{a} Z \cdot \text{EL222} \qquad\qquad \text{EL222} \xrightarrow{\lambda} \emptyset$$

$$\text{undifferentiated} \xrightarrow{u_s \cdot u(t) \cdot f(\text{EL222})} \text{differentiated}, \qquad (2)$$

where $u_s$ is the maximal single cell differentiation rate for given fixed light intensity, $u(t)$ is equal to one in the presence of light and equal to zero otherwise, $\lambda$ is the cells' growth rate, and $a$ is the rate at which protein bursts occur. Protein production bursts are of size $Z$ and assumed to be geometrically distributed with average burst size $b$, $Z \sim \text{Geo}(\frac{1}{b})$, as dictated by classical results for modeling stochastic gene expression [1]. We neglected delays and noise caused by stochastic production of recombinase from the optogenetic promoter for the sake of simplicity but also because previous modeling results for this optogenetic system suggested that cell-to-cell variability in the activity of the promoter is mostly a consequence of variability in transcription factor levels [2]. In the absence of light, $u(t) = 0$, all cells will remain undifferentiated and the EL222 distribution of the population converges approximately to a Gamma distribution (since the model is discrete it converges to a negative binomial distribution to be precise [3] but a Gamma distribution is more frequently used in the literature [1]). To be able to match this model to experimental fluorescence data, we assumed that molecule numbers and measured fluorescence are related via deterministic scaling with small zero-mean Gaussian measurement error

$$Y = s \cdot \text{EL222} + \varepsilon, \quad \text{where} \;\; \varepsilon \sim \mathcal{N}(0, \sigma^2). \tag{3}$$

The distribution of the data $Y$ is therefore obtained as the convolution of the scaled EL222 distribution with the distribution of $\varepsilon$. For all our results, the variance of the measurement error has been fixed to $\sigma^2 = 1.5$. Since this value is very small compared to typical fluorescence values, convolution of the EL222 distribution with the distribution of $\varepsilon$ has little practical consequences, except that it explains small tails into negative numbers of measured fluorescence distributions (see, for instance, Figure 2b in the main text). The scaling parameter $s$ is not identifiable from fluorescence measurements without any additional information on typical numbers of the transcription factor in cells. Furthermore, calculations with large transcription factor levels would be computationally infeasible without the use of sophisticated approximation techniques to reduce the number of states in the system's master equation [4] (in particular, for the composed model of the plasmid-based differentiation system). Therefore, we decided to fix $s = 5.06$a.u., which corresponds to very low transcription factor levels inside cells. We note that EL222 levels in cells matter primarily relative to the threshold of the differentiation Hill function $f(\text{EL222})$. Precise molecule numbers are of minor importance as long as the Hill function threshold is chosen in line with typical molecule numbers in the model. Furthermore, we note that the coefficient of variation of EL222 distributions generated by the model is fixed from the data. There is thus no increase in variability in the model due to small transcription factor numbers since any such increase is inherently corrected when the parameters of protein bursts are fixed. With $s$ (and $\lambda$) fixed, the model is reduced to burst frequency $a$ and average burst size $b$ as the two free parameters. Since these parameters are directly related to the mean and the coefficient of variation of the stationary negative binomial distribution of the model, they can be obtained from measured fluorescence distributions of cells growing in the dark. Concretely, we extracted mean and coefficient of variation of the EL222 distribution at the last measurement time point before light induction of the experiment in Figure 3a in the main paper. We find that the mean is equal to 50.6a.u. and the coefficient of variation is equal to 0.35. This allowed us to determine burst frequency and average size as $a = 0.0542\text{min}^{-1}$ and $b = 1.2426$ for the given value of $s$.

When light is applied to the population, $u(t) = 1$, the function $f(\text{EL222})$ determines which part of the cell population has a significant probability to differentiate. Since differentiation is caused by recombinase and recombinase has been omitted in the model, $f(\text{EL222})$ does not have a direct mechanistic interpretation. However, previous modeling results of the optogenetic system showed that cell-to-cell variability created by the system depends on whether its activity is modulated via light frequency or intensity [2]. This is explainable by a thresholding mechanism for the transcription factor in which, for low to intermediate light intensities, a fraction of the population does not have enough active EL222 to trigger significant activity of the optogenetic promoter. We therefore decided to model differentiation for our system with a Hill function $f(\text{EL222}) = \frac{\text{EL222}^{n_H}}{K_H^{n_H}+\text{EL222}^{n_H}}$ in line with the model of EL222-mediated gene expression in the work of Benzinger and colleagues [2]. We find that choosing a large Hill-exponent $n_H = 4$ and a Hill-threshold that is significantly larger than average EL222-levels, $K_H = 132/s$ (i.e. 132 in the fluorescence units displayed on x-axes of all distribution plots), coupled to a maximal single cell differentiation rate $u_s = 0.1\text{min}^{-1}$ leads to good agreement with observed population dynamics in Figure 3a in the main paper. The steepness of the Hill-function, coupled to a threshold that is far in the tail of the population's EL222 distribution, implies that the differentiation rate of the population is dependent on the rate at which fluctuations in EL222 replenish the tail of the EL222 distribution. We note that the time scale of fluctuations in constitutive expression of stable proteins (here of EL222) is determined by the cells' growth rate $\lambda$. Threshold-crossing "rates" for such proteins, however, have more complex dependencies on the protein production process and, in particular, depend on the size of fluctuations relative to the size of the threshold. Therefore, this model is capable of explaining slow population differentiation rates (e.g. that it takes approximately $20h$ for 90% of the population to differentiate in continuous presence of light, Figure 3b, bottom, in the main paper) without having to artificially impose a small value of $u_s$, which, in contrast, was necessary for the differentiation rate parameter ($u_{int} = 0.168\text{h}^{-1}$) of the simple deterministic model in Figure 1d in the main paper.

##### S.2.2 Mathematics of distribution dynamics in sub-populations.

In the absence of light, $u(t) = 0$, the stochastic model in Eq.(2) is a standard model of bursty gene expression and well-studied [1]. Its master equation is given by

$$\frac{d}{dt}p(x,t) = -(a+\lambda x)p(x,t) + \lambda(x+1)p(x+1,t) + a\sum_{y=1}^{x}\frac{1}{b}(1-\frac{1}{b})^{y-1}p(x-y,t), \tag{4}$$

where $p(x,t) := \mathbb{P}\left(X(t) = x \mid X(0) = x_0\right)$ and $X(t)$ is the number of proteins in a cell at time $t \geq 0$, $x \in \mathbb{N}_0$. To (approximately) calculate with this model, one can make use of a finite state projection [5] in which all protein production bursts that would lead the process to exit the truncated state space are instead redirected at a maximally allowed state $x_m$. $x_m$ needs to be chosen significantly larger than typical protein numbers for the finite state approximation to be accurate. Collecting all probabilities in a vector $\mathbf{p}(t) := [p(0,t) \ \cdots \ p(x_\text{m},t)]^T$, the master equation can be written in vector form as

$$\frac{d}{dt}\mathbf{p}(t) = \mathbf{A}\mathbf{p}(t), \tag{5}$$

where the entries of the generator matrix $\mathbf{A}$ are given as

$$\mathbf{A}_{x+1,y+1} = \begin{cases} -(a+\lambda x) & \text{if } y = x < x_m, \\ -\lambda x & \text{if } y = x = x_m, \\ \lambda(x+1) & \text{if } y = x+1, \\ \frac{a}{b}(1-\frac{1}{b})^{x-y-1} & \text{if } y < x < x_m, \\ \sum_{z=x-y}^{\infty}\frac{a}{b}(1-\frac{1}{b})^{z-1} & \text{if } y < x = x_m. \end{cases} \tag{6}$$

In the presence of light, $u(t) = 1$, this well-known model of gene expression is subject to state-dependent differentiation, which leaves the distribution of EL222 in the total population unaffected but creates complex dynamics in sub-populations. Possibly the most straightforward approach for obtaining distribution dynamics in sub-populations is to treat the differentiation state of the cell as an additional (pseudo-)species and to solve a new master equation with doubled state space size that basically consists of two coupled versions of Eq.(5). However, doubling the size of the state space, and the size of $\mathbf{A}$, implies that numerically solving the corresponding master equation may be significantly more difficult. Eq.(5) may, for instance, be solved via calculation of the matrix exponential of $\mathbf{A}$ but this becomes difficult if $\mathbf{A}$ is too large. On a more applied note, one can also realize that amounts of EL222 in differentiated cells are of no relevance for emerging population dynamics since these cells are already differentiated and will not differentiate again. In light of these considerations, it would be desirable to calculate with a small master equation that tracks only undifferentiated cells but, as opposed to Eq.(5), does so correctly in the presence of selective differentiation. Such an equation can be obtained by augmenting the bursty protein production model with an absorbing state, transitions to which represent a differentiation event in a cell. Denoting this new absorbing state by $D$, its probability by $p(D,t)$, and defining a new vector of all probabilities $\mathbf{p}_s(t) := [p(D,t)\ p(0,t)\ \cdots\ p(x_\mathrm{m},t)]^T$, we obtain the augmented master equation

$$\frac{d}{dt}\mathbf{p}_s(t) = \begin{bmatrix} 0 & \mathbf{c}_1 \\ \mathbf{0} & \mathbf{C} \end{bmatrix} \mathbf{p}_s(t), \tag{7}$$

where $\mathbf{c}_1 = u_s \cdot u(t) \cdot [f(0)\ f(1)\ \cdots f(x_m)]$ and $\mathbf{C}$ is the same as $\mathbf{A}$ in Eq.(6) except that the outflow terms due to differentiation in $\mathbf{c}_1$ are subtracted from the diagonal of $\mathbf{A}$.

To obtain the dynamics of EL222 distributions in undifferentiated cells, we can define the conditional probabilities $p_c(x,t) := \mathbb{P}\left(X(t) = x \mid X(t) \neq D, X(0) = x_0\right)$ for a cell to have x proteins at time $t$ given that it has not differentiated yet. Collecting conditional probabilities in a vector $\mathbf{p}_c(t) := [p_c(0,t)\ \cdots\ p_c(x_\mathrm{m},t)]^T$, it is straightforward to show that the EL222 distribution in undifferentiated cells follows the non-linear equation

$$\frac{d}{dt}\mathbf{p}_c(t) = \mathbf{C}\mathbf{p}_c(t) + \mathbf{p}_c(t) \cdot (\mathbf{c}_1 \mathbf{p}_c(t)). \tag{8}$$

When light is continuously maintained, $u(t) = 1$, $\mathbf{p}_c(t)$ converges to a quasi-stationary distribution, $\mathbf{p}_\mathrm{QSD}$, which can be obtained as the normalized eigenvector of $\mathbf{C}$ corresponding to the largest eigenvalue [6]. Correspondingly, the (per cell) population differentiation rate converges to the negative of this eigenvalue (the largest eigenvalue is always negative). In the absence of light, $u(t) = 0$, it holds that $\mathbf{c}_1 = \mathbf{0}$ and $\mathbf{C} = \mathbf{A}$, which implies that the EL222 distribution follows the standard master equation of the bursty protein production model and reverts back to its well-known stationary negative binomial distribution.

##### S.2.3 Selection effects for the integrated differentiation system.

To test our mathematical results experimentally, we exposed cells to continuous light and measured the resulting dynamics of the EL222 distribution of the undifferentiated cell population. Since the continuous presence of light eventually leads to differentiation of the entire population, we can only reliably quantify EL222 distributions in undifferentiated cells at early enough time points with sufficiently many undifferentiated cells. Figure S.4 shows that both model and data distributions start to shift after the application of light. The bottom panels ($11-15$h after light induction) indicate that distributions do not significantly shift further after some time. According to the model, these distributions represent the quasi-stationary condition that balances selective differentiation of cells with high EL222 levels and replenishment of the original distribution due to the stochastic protein production process.

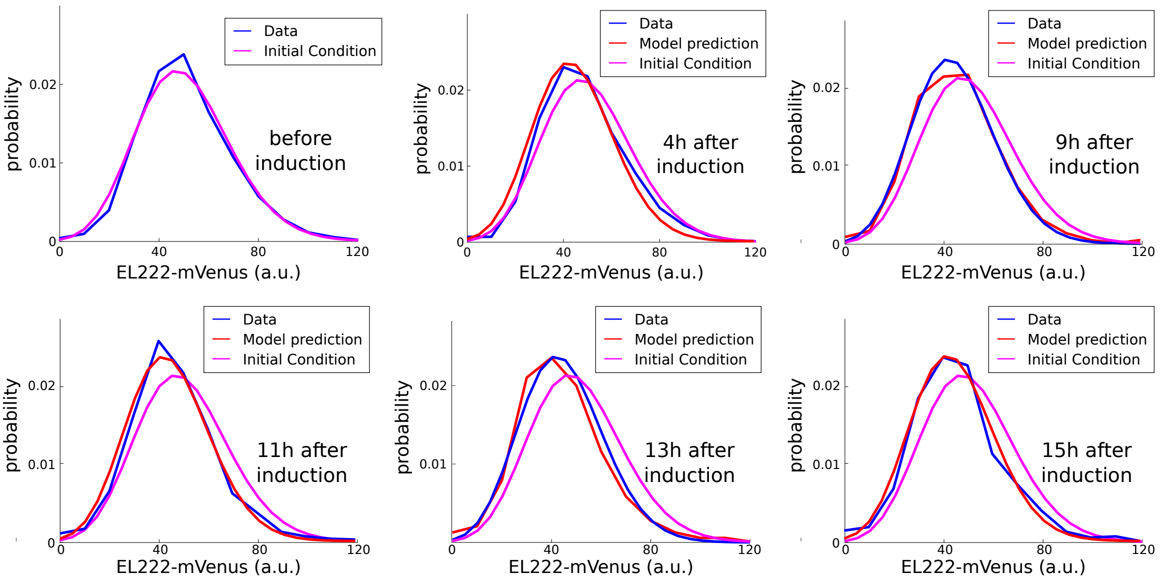


Figure S.4: **EL222 distribution dynamics in continuous light**. The initial condition of the model (magenta in all panels) is the negative binomial distribution of the bursty protein production model. When light is applied, cells with high EL222 levels differentiate and EL222 distributions of cells that remain undifferentiated shift to lower levels according to both model (red) and data (blue). According to the model, a quasi-stationary condition is reached some hours after the start of light induction.

In addition to the experiment shown in Figure S.4, we performed another experiment where light is applied for some time but then removed again. However, given the only very small shift in distributions, we could not clearly establish that the EL222 distribution of undifferentiated cells reverts back to the initial condition in the dark for the integrated system. We refer the reader to the plasmid-based version of the system for a demonstration of this result (e.g. Figure 5 in the main text).

### S.3 Plasmid dynamics and population net growth rate.

#### S.3.1 Construction of the model.

The population dynamics of cells that have or have not lost the plasmid can be described by the following ordinary differential equations

$$\begin{aligned} \frac{d}{dt}n &= \lambda n - ln \\ \frac{d}{dt}n_0 &= (\lambda - \mu)n_0 + ln, \end{aligned} \tag{9}$$

where $n$ is the (expected) number of cells that have not lost the plasmid, $n_0$ is the number of cells that have lost the plasmid, $\lambda = 0.0067\ \text{min}^{-1}$ is the single-cell growth rate (corresponding to a division time of 103min), $l$ is the plasmid loss rate, and $\mu$ represents a removal rate of cells that have lost the plasmid. Taking a single-cell perspective, plasmid loss is a cellular event and it is a priori unclear if it can be appropriately described by a single rate parameter $l$ since such a description implicitly carries the assumption that the waiting time for the event to occur follows an exponential distribution. However, one can show that the simple population dynamics model Eq.(9) can be derived from a mechanistic representation of single-cell plasmid copy number fluctuations where the plasmid loss rate emerges as the difference between dilution and replication rate, $l = \lambda - a_p$ (see the following Section S.3.2).

In non-selective media, it holds that $\mu = 0$, the total population grows at a rate $\lambda$, and the ratio $\frac{n}{n_0+n}$ will converge exponentially to zero at a rate that is determined by the plasmid loss rate $l$. Similarly, average plasmid copy numbers of cells in the population decrease exponentially at rate $l$ when cells are switched from selective to non-selective media (see the following Section S.3.2). Assuming that the total rate of protein production of a cell is linear in the number of plasmids, then implies that expression levels of constitutive proteins approximately decay exponentially at rate $l$ since, from a mechanistic perspective, the plasmid loss rate must be smaller than the growth rate $\lambda$ and therefore the slowest time scale that determines changes in protein levels. Since gene expression levels can readily be measured, we can experimentally quantify $l = 10^{-3}$ min$^{-1}$ by switching cells from selective to non-selective media and observing the decay rate of the mean of a constitutively expressed protein (Figure 4a in the main paper). From this result, the plasmid replication rate can be derived as $a_p = 0.0057$ min$^{-1}$ $= 0.85\lambda$, i.e. that each plasmid is only replicated successfully with a probability of 0.85.

In selective media, $\mu$ will take a strictly positive value and must be sufficiently large compared to $l$ to ensure that cells with plasmids can be maintained in the population. If this is the case, straightforward manipulation of Eq.(9) shows that the fraction of cells with plasmids converges to $\frac{n}{n_0+n} = \frac{\mu-l}{\mu}$ and the net population growth rate to $\lambda_{\text{select}} = \lambda - l = a_p$. The seemingly counterintuitive result that the growth rate in selective media does not depend on $\mu$, and is instead equal to the plasmid replication rate, is due to the fact that reduced growth rates for cells without plasmids imply that the fraction of these cells will be correspondingly smaller in stationary growth conditions. This implies that growth rate measurements in selective media cannot be used to determine $\mu$ but instead provide an independent measurement of $a_p$. We find that selective media reduces the population growth rate by about 15% (Figure 4b in the main paper), which is in agreement with the previous finding that the plasmid replication rate must be 15% smaller than $\lambda$.

With the value of $l$ (double-) confirmed, we set out to determine the value of $\mu$ by measuring the fraction of cells without plasmids in selective media, $\frac{n_0}{n_0+n}$. To this end, we performed a colony counting experiment (see also Section S.7.1). Briefly, we cultured cells harboring the integrated system and cells harboring the plasmid-based system to exponential phase in selective media in triplicates. We then did serial dilutions of the cultures and plated them on selective and non-selective media. After 48h of incubation at 30°C in the dark, we counted the number of colonies on each plate manually (Figure 4a in the main text). We found that, while there was no significant difference in the number of colony forming units (CFUs) at different dilutions between selective and non-selective media for the strain with the integrated system, the number of CFUs for the strain with the plasmid-based system was always lower compared to the strain with the integrated system. On average, upon normalization with the CFUs observed in non-selective media at the given dilution, we observed a significant decrease in the number of CFUs on selective media plates for the strain with the plasmid-based system (Figure 4a in the main text). Comparing the values, we estimated that around a third ($32.83\% \pm 10.16\%$ (mean $\pm$ s.d.)) of cells do not contain plasmids during exponential growth.

According to the modeling and experimental results presented in this section, our cells carry on average around 4 plasmids during stationary growth in selective media. Average plasmid copy numbers of cells carrying plasmids similar to the ones used in the study have been reported in the past via bulk measurement methods like DNA extraction of the whole population followed by qPCR [7, 8, 9]. There is significant variability in the numbers reported in different studies which is understandable given that each study used different plasmid backbones (the backbones in Gnügge et al. [9] are closest to ours). In Gnügge et al. [9], an approach with two constitutively expressed fluorescence reporters, one integrated in the genome while the other destabilized and expressed from a plasmid, was employed to indirectly estimate the distribution of 2-micron plasmid copy numbers in the population via flow cytometry. The authors observed that a significant fraction of the population lacked plasmids (45%). The fluorescence distribution of the destabilized constitutively expressed gene resemble the mVenus fluorescence distribution of our system, except that peaks in the distribution, corresponding to different plasmid copy numbers according to Gnügge et al., are smeared out in our data since we did not use a destabilized reporter.

##### S.3.2 Mathematics of the single cell plasmid model in growing populations.

We represent the dynamics of plasmid copy numbers in growing cell populations by a one-dimensional replication-dilution process coupled to a removal process for cells that have lost the plasmid

$$\begin{aligned} &P \xrightarrow{a_p} P + P \qquad\qquad P \xrightarrow{\lambda} \emptyset \\ &\text{cell} \xrightarrow{\mu \cdot \mathbb{1}_{\{P=0\}}} \text{removed}, \end{aligned} \tag{10}$$

where $a_p$ is the plasmid replication rate, $\lambda$ is the growth rate of cells and thus the rate at which plasmids are diluted, and $\mu$ is the rate at which cells that have lost the plasmid are removed from the population. In non-selective media, $\mu = 0$, this process is governed by the following master equation:

$$\frac{d}{dt}p(x,t) = -(a_p + \lambda)xp(x,t) + a_p(x-1)p(x-1,t) + \lambda(x+1)p(x+1,t), \tag{11}$$

where $p(x,t) := \mathbb{P}\left(X(t) = x \mid X(0) = x_0\right)$ and $X(t)$ is the number of plasmids in a cell at time $t \geq 0$, $x \in \mathbb{N}_0$. Collecting the probabilities of all states in a vector, $\mathbf{p}(t) := [p(0,t) \;\cdots\; p(x_\mathrm{m},t)]^T$, and projecting on a finite state space it is well known that the master equation can be written in vector form as

$$\frac{d}{dt}\mathbf{p}(t) = \mathbf{A}\mathbf{p}(t) \tag{12}$$

$$\mathbf{A} = \begin{bmatrix} 0 & \lambda & 0 & \cdots & \cdots & \cdots & \cdots & 0 \\ 0 & -(\lambda + a_p) & 2\lambda & 0 & \cdots & \cdots & \cdots & 0 \\ 0 & a_p & -2(\lambda + a_p) & 3\lambda & 0 & \cdots & \cdots & 0 \\ 0 & 0 & 2a_p & -3(\lambda + a_p) & 4\lambda & 0 & \cdots & 0 \\ \vdots & \vdots & & & \ddots & & & \vdots \\ 0 & 0 & \cdots & 0 & (x_\mathrm{m}-3)a_p & -(x_\mathrm{m}-2)(\lambda + a_p) & (x_\mathrm{m}-1)\lambda & 0 \\ 0 & 0 & \cdots & \cdots & 0 & (x_\mathrm{m}-2)a_p & -(x_\mathrm{m}-1)(\lambda + a_p) & x_\mathrm{m}\lambda \\ 0 & 0 & \cdots & \cdots & \cdots & 0 & (x_\mathrm{m}-1)a_p & -x_\mathrm{m}\lambda \end{bmatrix}.$$

For this finite state version to be an accurate approximation of the infinite state Markov chain, $x_\mathrm{m}$ needs to be chosen large enough for it to be unlikely that the process ever reaches such large plasmid copy numbers such that the omission of outflow and inflow terms beyond the truncation (in the last row) does not lead to noticeable changes in the solution.

Due to the absorbing state at zero, the stationary distribution of this Markov chain has probability one in the state zero, corresponding to all cells in a population having lost the plasmid, or alternatively to any single cell having lost the plasmid with probability one. When cells are grown in selective media, however, cells that lose the plasmid are gradually removed from the population, for instance because they die. To obtain an equation that describes plasmid copy number distributions for populations growing in selective media, we can add a new absorbing state $D$ for dead cells that is reachable from $x = 0$ with a rate $\mu$ that represents the death of cells without plasmids. Defining $p(D,t) := \mathbb{P}\left(X(t) = D \mid X(0) = x_0\right)$ and $\mathbf{p}_s(t) := \left[p(D,t)\ p(0,t)\ \cdots\ p(x_{\mathrm{m}},t)\right]^T$, we can write a new master equation for cells in selective media as

$$\frac{d}{dt}\mathbf{p}_s(t) = \begin{bmatrix} 0 & \mathbf{c}_1 \\ \mathbf{0} & \mathbf{C} \end{bmatrix} \mathbf{p}_s(t), \tag{13}$$

where $\mathbf{c}_1 = [\mu\ 0\ \ldots 0]$ and $\mathbf{C}$ is the same matrix as $\mathbf{A}$ in Eq.(12) except that the zero in the $(1,1)$-entry of $\mathbf{A}$ is replaced by $-\mu$.

Eq.(13) is a useful tool as it allows us to define the conditional probabilities $p_c(x,t) := \mathbb{P}\left(X(t) = x \mid X(t) \neq D, X(0) = x_0\right)$ for a cell to have x plasmids at time $t$ given that it has not died yet. Collecting conditional probabilities in a vector $\mathbf{p}_c(t) := [p_c(0,t)\ \cdots\ p_c(x_{\mathrm{m}},t)]^T$, it is straightforward to show that the plasmid copy number distribution for all cells that are part of the population at time $t$ follows the same type of non-linear equation as obtained previously (Eq.(8)) for EL222 distributions in undifferentiated cells:

$$\frac{d}{dt}\mathbf{p}_c(t) = \mathbf{C}\mathbf{p}_c(t) + \mathbf{p}_c(t) \cdot \left(\mathbf{c}_1 \mathbf{p}_c(t)\right). \tag{14}$$

When growth conditions do not change, $\mathbf{p}_c(t)$ converges to a quasi-stationary distribution, $\mathbf{p}_{\mathrm{QSD}}$, which can be obtained as the normalized eigenvector of $\mathbf{C}$ corresponding to the largest eigenvalue. While the above derivation may seem complex, we note that it is highly useful: plasmid copy numbers are typically measured in populations growing in selective media and Eq.(14) is necessary to correctly interpret resulting data. For instance, average plasmid copy numbers can be measured by qPCR and this average can be compared to the mean of $\mathbf{p}_{\mathrm{QSD}}$. Furthermore, Eq.(14) allows us to consolidate the mechanistic single cell perspective of plasmid loss events with the population perspective of a measurable plasmid loss rate (as in Eq.(9)). When populations are grown in selective media, their plasmid copy number distribution equilibrates to the quasi-stationary $\mathbf{p}_{\mathrm{QSD}}$. Switching to non-selective media then frees up only one-dimensional dynamics and the plasmid copy number distribution will gradually shift towards $x = 0$ while leaving the ratios of probabilities of all other states unchanged. This implies that there exists a single time scale, and a single rate, that emerges as a plasmid loss rate at the population scale. This rate can be obtained as the negative of the largest eigenvalue of $\mathbf{C}$. For the simple replication-dilution model that we used here, we find that the negative of the largest eigenvalue of $\mathbf{C}$ is equal to $\lambda - a_p$, which is in line with the intuitive expectation that a plasmid loss rate that emerges at the population scale should be determined by the difference of replication and growth rate.

#### S.4 Composed model of plasmid copy numbers and differentiation dynamics.

##### S.4.1 Calculating with the composed model

Master equation and generator matrix of the composed model can be obtained akin to the component models. To be able to compare distributions of both undifferentiated and differentiated cells to data (Figure 5 in the main paper), as opposed to Section S.2.2, we have decided to explicitly track also differentiated cells even though their statistics are not needed to determine differentiation dynamics at the population scale. This comes at the cost of a doubled state space size, as already stated in Section S.2.2, and leads to a normal linear master master equation. Eventually, we still require a non-linear master equation of the type

$$\frac{d}{dt}\mathbf{p}_c(t) = \mathbf{C}\mathbf{p}_c(t) + \mathbf{p}_c(t) \cdot (\mathbf{c}_1 \mathbf{p}_c(t)) \tag{15}$$

for the composed model since the plasmid part of the model requires conditioning on cells not being removed due to plasmid loss, as described in detail in Section S.3.2. The generator matrix of the composed model is too unwieldy to reproduce here since the model is 3-dimensional (EL222, plasmids, differentiation status). In our code, it is constructed algorithmically with for-loops running through all dimensions and all possible state changes coupled to an alignment of the 3-dimensional (finitely truncated) state space in a single dimension to obtain a vector form of the master equation. We tested accuracy of finite state projections of varying sizes and eventually settled for a truncation $251 \cdot 31 \cdot 2$ that leads to a generator matrix of size $15562 \times 15562$. For all calculations where this is possible, we dropped the differentiated population from the model and calculated with a generator matrix of only half the size. For instance, the initial condition of undifferentiated cells in the model (EL222 and plasmid distributions in the dark, Figure 5 in the main paper and Figure S.6) are determined as eigenvectors of a matrix $\mathbf{C}$ that is only $7781 \times 7781$.

For the plasmid-based system, there is a small fraction of cells that are already differentiated before light is applied. For the modeling, we assumed that these cells differentiated long before the start of the experiment such that their EL222 and plasmid distributions have already equilibrated back to the same distribution as that of undifferentiated cells. We note, however, that it is also possible that these cells recombined due to ambient light contamination when the cells are handled for loading into the bioreactor platform. This constitutes a possible source of mismatch between the composed model and experimental data.

To obtain transient distribution dynamics in response to light, we dropped the non-linear terms in Eq.[15] from the model and removed the absorbing state corresponding to cells that are removed due to plasmid loss while keeping the outflow terms due to cell removal on the diagonal of $\mathbf{C}$. This leads to a linear version of the equation that can be solved, for instance, via matrix exponentiation [10] but that does not preserve the total probability mass. We have shown previously, however, that renormalizing the resulting solution to a probability distribution allows one to obtain the solution of Eq.(15) without the need to solve a non-linear equation [4].

##### S.4.2 Additional results for the plasmid-based differentiation system

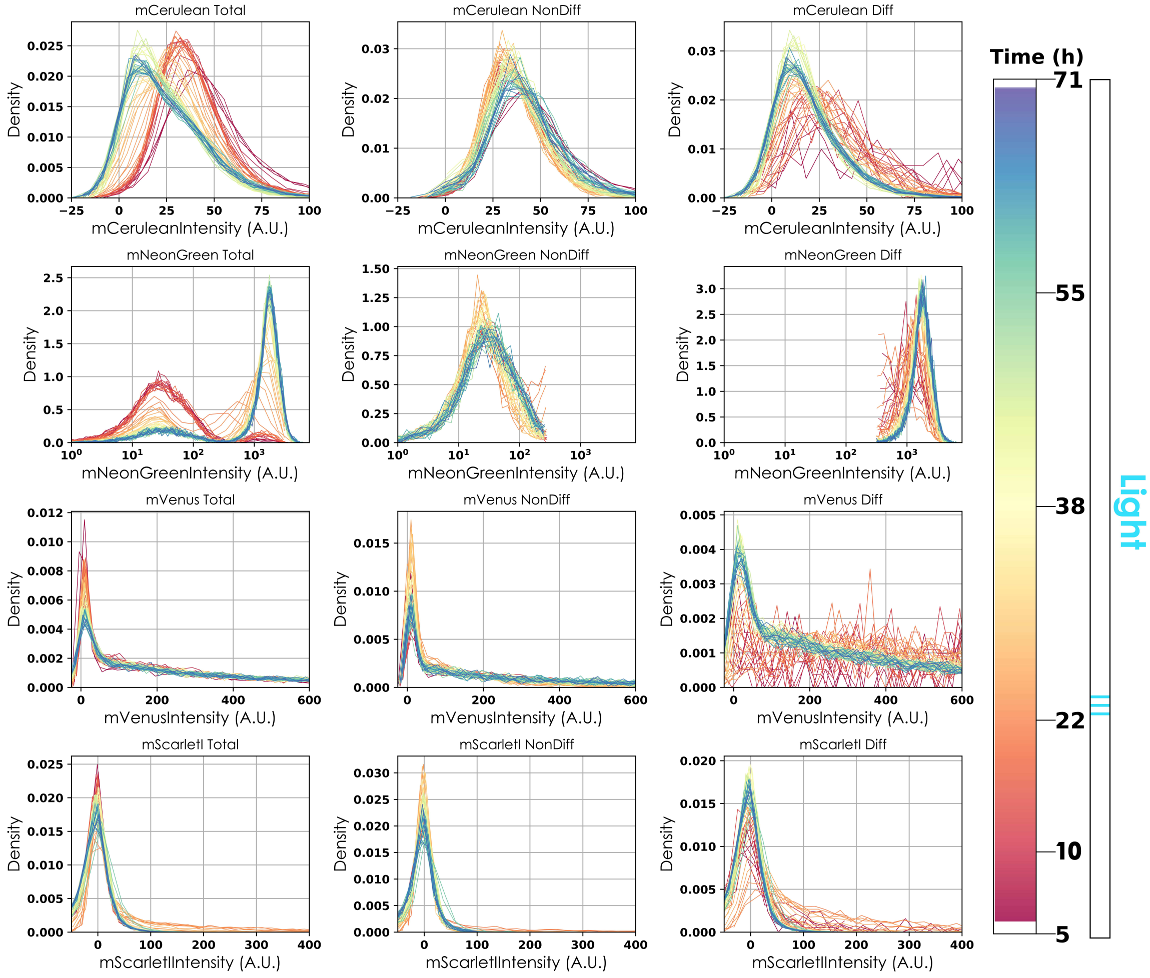


Figure S.5: **Distribution dynamics of the four fluorescent reporter proteins for the experiment in Figure 5 in the main paper.** Color coding indicates the time of the experiment. Rows correspond to the different fluorescent reporters, columns to different sub-populations (left: all cells, middle: undifferentiated cells, right: differentiated cells) after classification based on mNeonGreen abundance. We note that due to changing sizes of sub-populations, fluorescence distributions need to be extracted from varying (sometimes low) cell numbers.

In the main text, we have shown that the composed model leads to accurate predictions of differentiation dynamics at the population scale as well as good agreement of predicted dynamics of EL222 distributions with experimental data. Full time-varying distribution data of all fluorescent reporters for this experiment is displayed in Figure S.5.

Dynamics of plasmid copy number distributions in sub-populations cannot be measured directly in experiments. They are nonetheless interesting and can readily be obtained from the model. Figure

S.6 shows model-predicted plasmid copy number distributions for the experiment and time points displayed in Figure 5 in the main paper. According to the model, cells contain on average around 4 plasmids. After exposure to light, the average plasmid copy number in differentiated cells increases to more than 6 while cells that remain undifferentiated have on average only slightly more than two plasmid. When cells are maintained in the dark, plasmid copy number distributions of both sub-populations converge back to their initial condition.

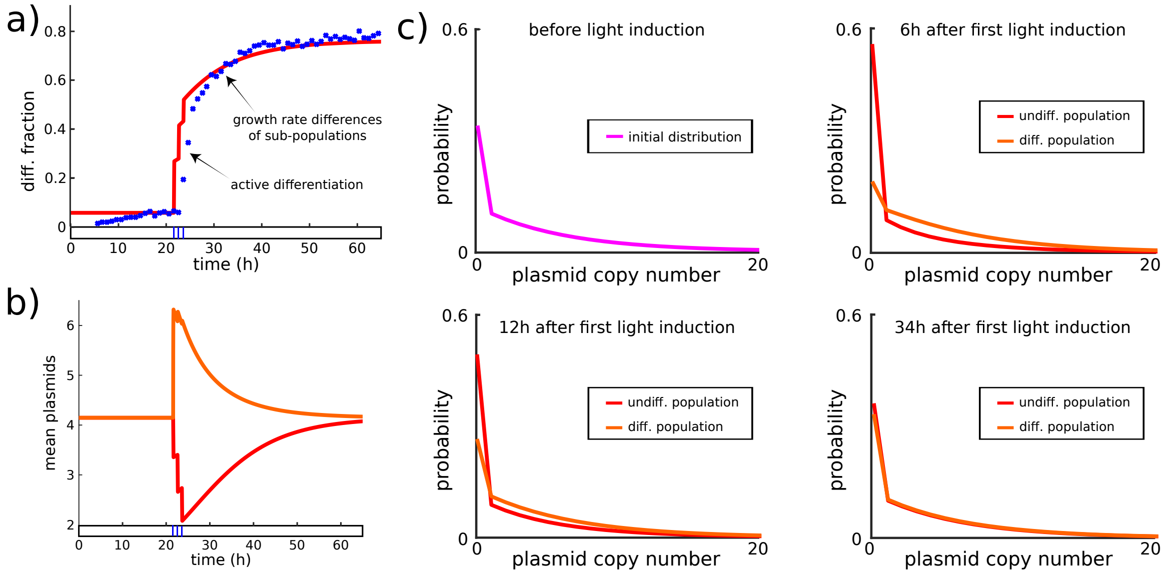


Figure S.6: **Additional results for the experiment in Figure 5 in the main paper. (a)** Emerging population dynamics, same as Figure 5 in the main paper. **(b)** Dynamics of average plasmid copy numbers in sub-populations. **(c)** Dynamics of sub-population plasmid copy number distributions. Color coding and time points are the same as in Figure 5 in the main paper.

Differing plasmid copy number distributions lead to different removal rates of cells from sub-populations due to plasmid loss. Consequently, effective sub-population growth rates in selective media will be transiently different. Model-predicted sub-population growth rates are shown in Figure S.7a. Differing sub-population growth rates imply that the differentiated population fraction continues to increase in the dark even though there is no more active differentiation (Figure 5 in the main paper and Figure S.6a). Sub-population growth rates are not experimentally measurable since all cells grow together and the total population growth rate remains constant at all times according to both model and data. The continuing increase in the differentiated fraction in the dark is, however, experimentally observable and agrees well with predictions of the model.

Figure S.7 also shows that multiple light pulses lead to a slightly reduced population differentiation rate for subsequent light pulses compared to the first. This is a consequence of selective differentiation of cells with high EL222 levels in the first stimulus and plasmid mixing dynamics that are too slow to replenish the sub-population plasmid copy number distribution in between pulses. Selective differentiation therefore creates a transient memory of the light stimulus that is retained for a time that depends on how plasmid copy numbers fluctuate in cells.

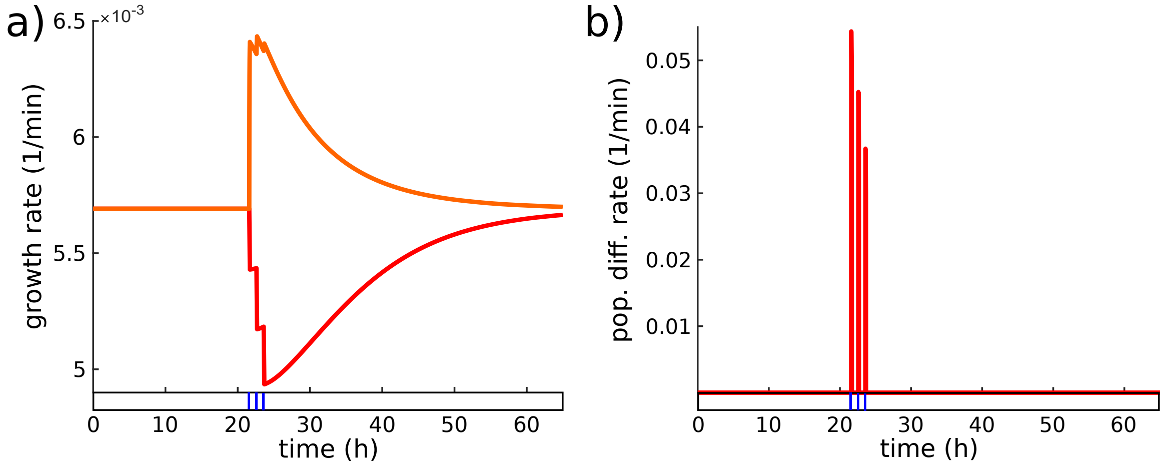


Figure S.7: **Additional results for the experiment in Figure 5 in the main paper. (a)** Sub-population growth rates according to the model of undifferentiated (red) and differentiated (orange) cells. **(b)** Population differentiation rate according to the model.

##### S.4.3 Differention dynamics for varying light sequences

To test the plasmid-based differentiation system and to determine how well other experiments can be predicted by the composed model, we exposed cells to light input sequences different from the one shown in Figure 5 in the main paper. To start, we chose a light sequence with light pulses that are of the same duration (5min) as in Figure 5 in the main paper but that are applied further apart from each other (4h between subsequent pulses instead of 1h). Overall, we applied 5 such light pulses as opposed to the three pulses in the experiment of Figure 5 in the main paper. Full time-varying distribution data of all fluorescent reporters for this experiment is displayed in Figure S.8. We find that the experimentally measured differentiated fraction grows more slowly compared to Figure 5 in the main paper (Figure S.9a). As for previous results, EL222 distributions in undifferentiated cells shift to lower levels in response to light (Figure S.9b,d). Both differentiation dynamics and dynamics of sub-population EL222 distributions are well predicted by the composed model. A notable difference to the experiment in Figure 5 in the main paper is that 10h after the last light signal there is almost no further increase in the differentiated fraction. According to the model, this is a consequence of the increased spacing between subsequent pulses that allows plasmid copy numbers and sub-population growth rates more time to relax back to their initial conditions (Figure S.9c).

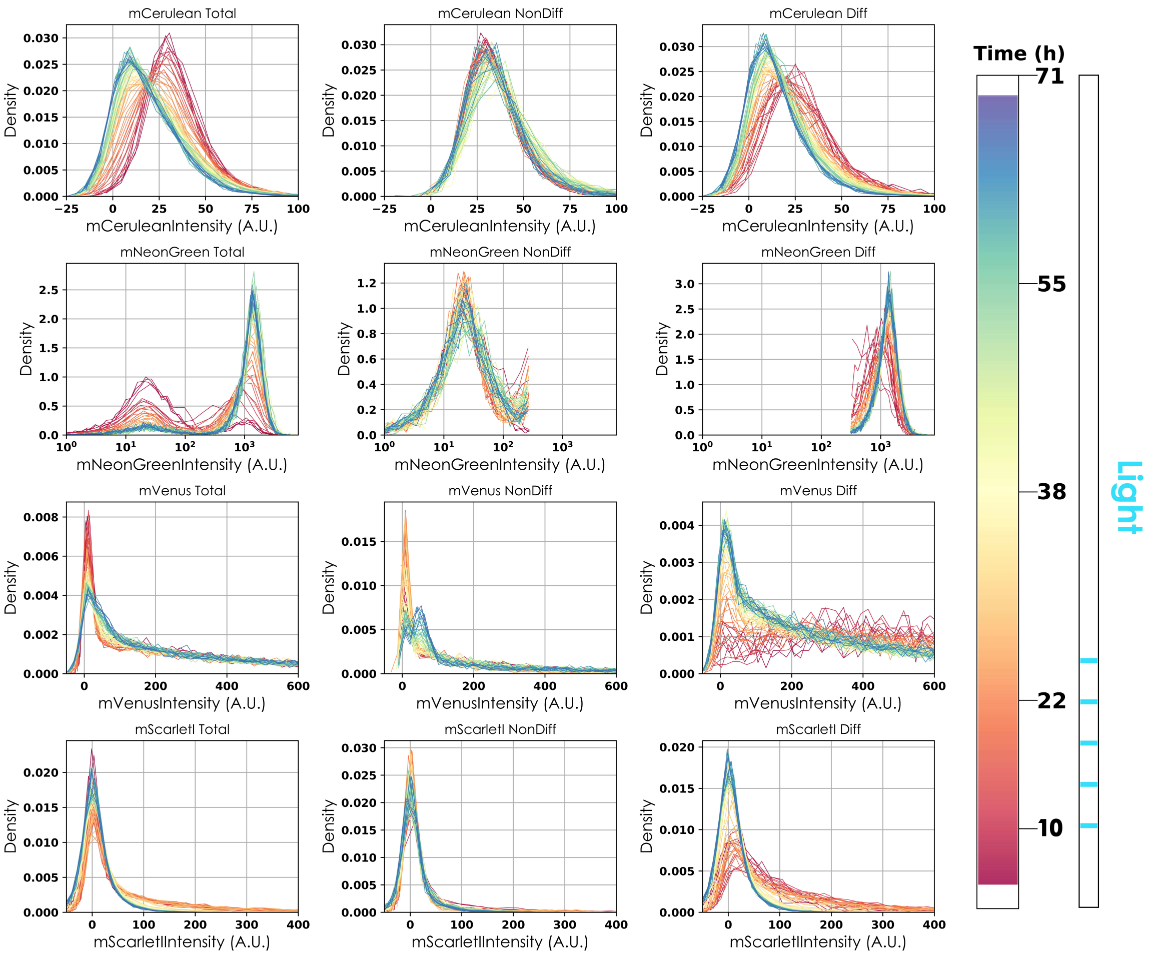


Figure S.8: **Distribution dynamics of the four fluorescent reporter proteins for the experiment with five 5min light pulses** (see also Figure S.9). Color coding indicates the time of the experiment. Rows correspond to the different fluorescent reporters, columns to different sub-populations (left: all cells, middle: undifferentiated cells, right: differentiated cells) after classification based on mNeonGreen abundance. We note that due to changing sizes of sub-populations, fluorescence distributions need to be extracted from varying (sometimes low) cell numbers.

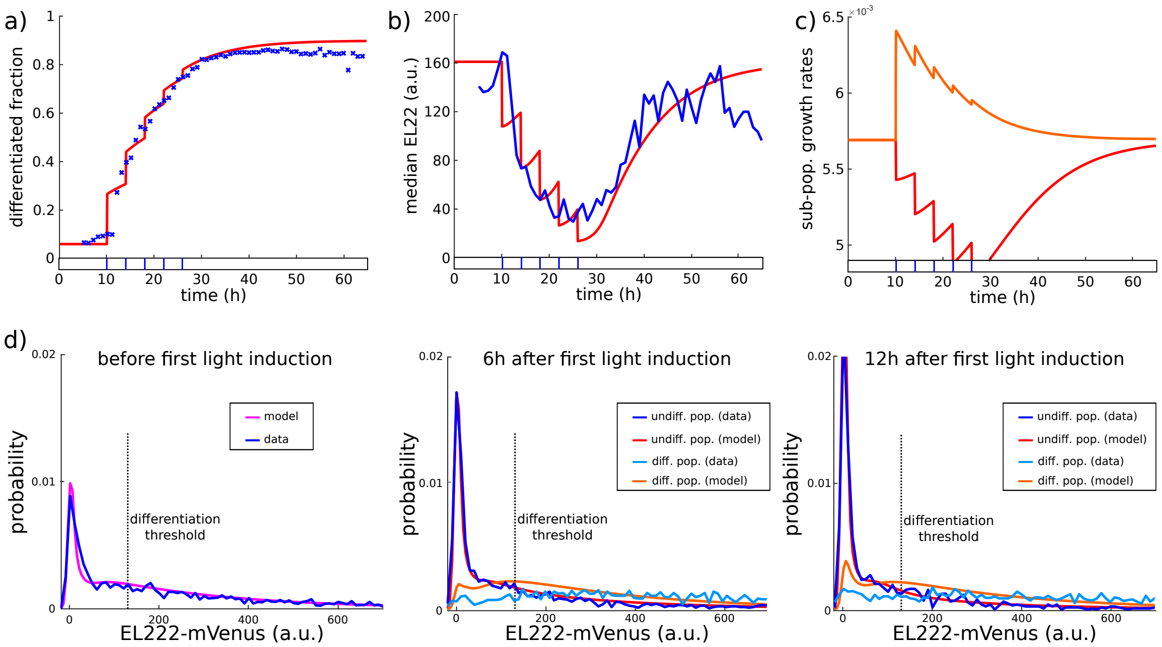


Figure S.9: **Response of the plasmid-based system for another light sequence. (a)** Emerging population dynamics according the single cell model (red) for the light input shown at bottom are compared to experimental data (blue). **(b)** Dynamics of the median of EL222 distributions according to the model (red) and data (blue). **(c)** Sub-population growth rates of undifferentiated (red) and differentiated (orange) cells according to the model. **(d)** Initial EL222 distribution in the dark (left) and distributions shifts after light induction (middle and right).

Subsequently, we applied continuous light to cells and found that the entire population recombines fairly quickly despite the presence of $\sim 33\%$ cells that have mVenus levels close to zero and seem to have lost the plasmid (Figure S.10). This is not a contradiction since populations grow in selective media and, with the help of the model, we established that such cells are continuously removed from the population while at the same time new cells without plasmids emerge due to plasmid loss events. New plasmid loss events may equally well happen in cells that are already differentiated.

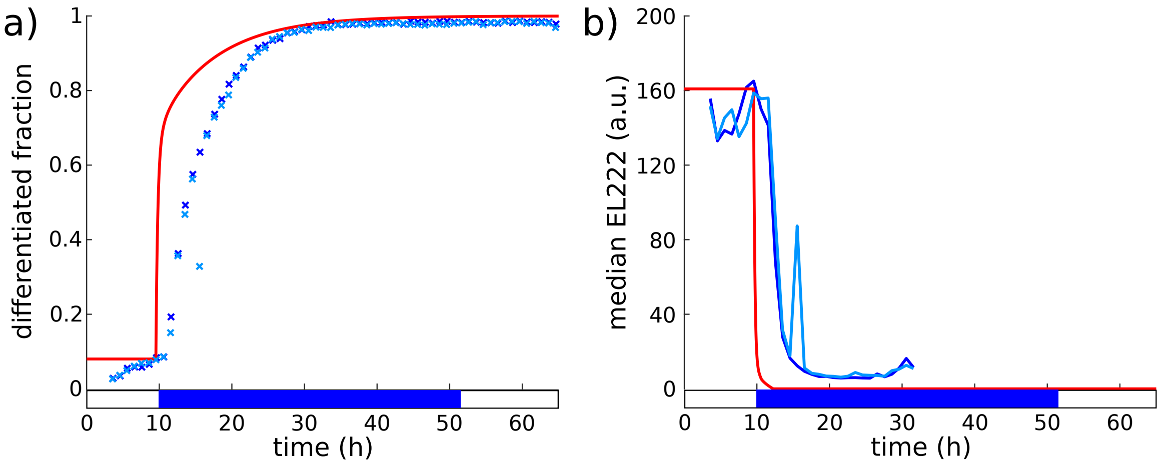


Figure S.10: **Plasmid-based system in continuous light (a)** Differentiation dynamics obtained from two replicates (shades of blue) are compared to predictions of the composed model (red). Replicates agree perfectly but there are differences to the model in early transients since the model neglects delays due to recombinase production and experimental detection of differentiated cells (needs enough mNeonGreen production and maturation). **(b)** Dynamics of EL222 medians for the two replicates. Under continuous light, the median drops to almost zero since at all time points more than half of the undifferentiated cells have no plasmids (according to the model). At later time points ($>$ 20h after the start of light induction) most cells are recombined and the median of undifferentiated cells cannot be reliably computed.

#### S.5 Experimental replicates

Since we have made extensive use of particular experimental results for constructing and testing the composed model, it is important to verify that these results are repeatable. First things first, a certain degree of variability in experiments on our bioreactor platform cannot be avoided. In particular, the different reactors of the platform are each equipped with their individual LEDs. In principle, all LEDs are identical but a certain degree of variability in light intensity between LEDs for the same setting cannot be avoided, in particular when the intensity is set to be low. To reduce the consequences of such variability, we have only used a fixed fairly large intensity setting for all experiments in the paper. Nevertheless, the same intensity setting may lead to slightly more or less differentiation in different experiments. We note, however, that we only ever observed minor differences. Furthermore, LED variability creates only variations in the system's input that does not alter core dynamical processes in any way. Figures S.11a shows three replicates of the experiment with the integrated differentiation system in Figure 3a in the main paper that was used for constructing the model. We find that these experiments are in good agreement and show very similar dynamics of the differentiated fraction in response to light except that a slightly smaller differentiated fraction emerged for one of the experiments (presumably due to LED variability). Two replicates of the continuous-light experiment in Figure 3b in the main paper show also very good agreement (Figures S.11b).

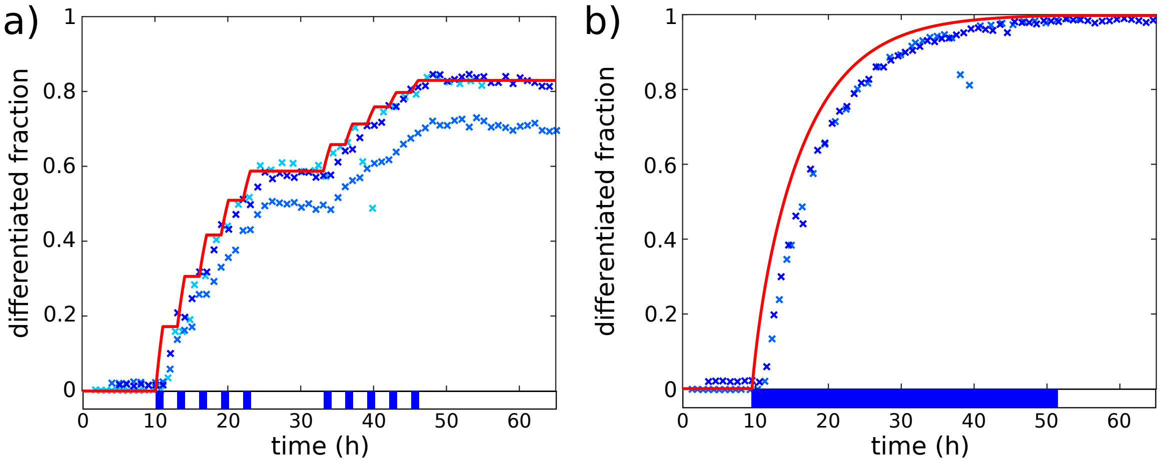


Figure S.11: **Replicates for the integrated differentiation system.** **(a)** Dynamics of the differentiated fraction in three replicates of the experiment in Figure 3a in the main paper. Absolute experiment time on the x-axis of some of the replicates has been shifted to align the timing of the light pulses in the different experiments. **(b)** Dynamics of the differentiated fraction in two replicates of the experiment in Figure 3b in the main paper.

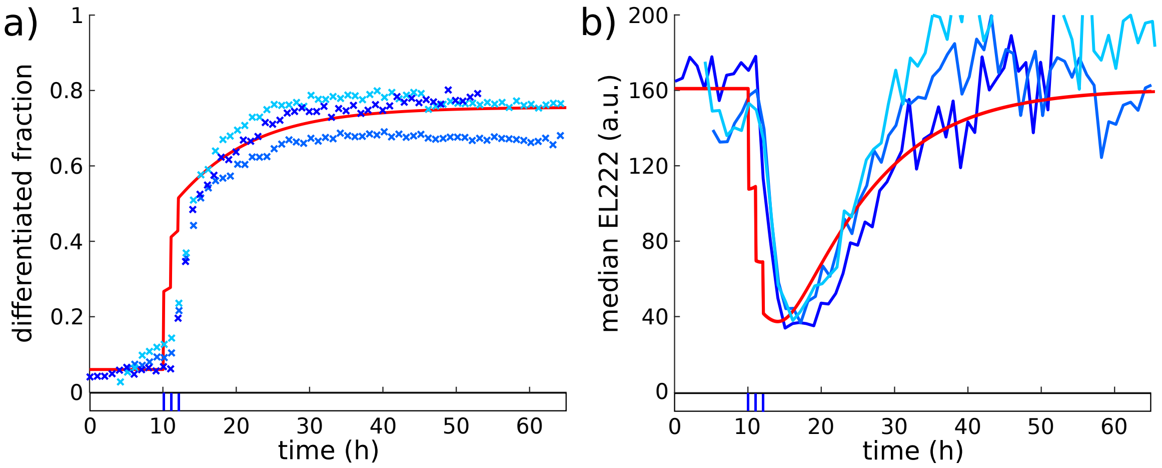


Figure S.12: **Replicates for the plasmid-based differentiation system.** **(a)** Dynamics of the differentiated fraction in three replicates of the experiment in Figure 5 in the main paper. **(b)** Median EL222 dynamics for the same experiments as in panel a. In both panels, absolute experiment time on the x-axis of some of the replicates has been shifted to align the timing of the light pulses in the different experiments.

For the plasmid-based system, an additional difficulty is that the system is more light sensitive than the integrated system (less light is required to differentiate cells with many plasmids). As a consequence, it is hard to avoid that some cells differentiate due to ambient light contamination when cells are loaded into the bioreactor platform. This may lead to non-zero differentiated fractions at the start of the experiment and slightly reduced levels of EL222 in undifferentiated cells at early time points prior to blue light exposure. Figure S.12 shows three replicates of the experiment in Figure 5 in the main paper. Overall, replicates agree very well in the dynamics of the differentiated fraction (Figures S.12a) and the shift in EL222 levels in undifferentiated cells (Figures S.12b) with some small differences being present due to the aforementioned LED variability and ambient light contamination.

#### S.6 Repeated light pulses

In the main paper, we used repeated light pulses to regulate constitutive gene expression levels in the undifferentiated sub-population to constant lowered levels. Figure S.13 supplements the results in Figure 6 in the main paper and displays the full time-varying distribution data of all fluorescent reporters for the experiment in which cells have been exposed to 2min of light every 4h. Particular attention should be given to the third row in Figure S.13 that provides EL222 distribution dynamics. The middle panel shows EL222 distributions in undifferentiated cells. The shift from red to orange and yellow colors corresponds to the shift from the magenta to the blue distribution in Figure 6,c in the main paper. The distribution then remains approximately invariant for tens of hours (yellow and orange). Towards the end of the experiment (green and blue colors), there seems to be a reduced number of cells that contain no plasmids (reduced peak at zero) and an increased number of cells with approximately one plasmid (despite the fact that the median of the distribution stays more or less constant (Figure 6,b in the main paper)). This change in EL222 distributions manifests itself also in the distribution of the entire population (third row, left panel). We must therefore conclude that plasmid regulation is modified towards the end of the experiment in some unknown way despite the fact the the media of the experiment has not been changed.

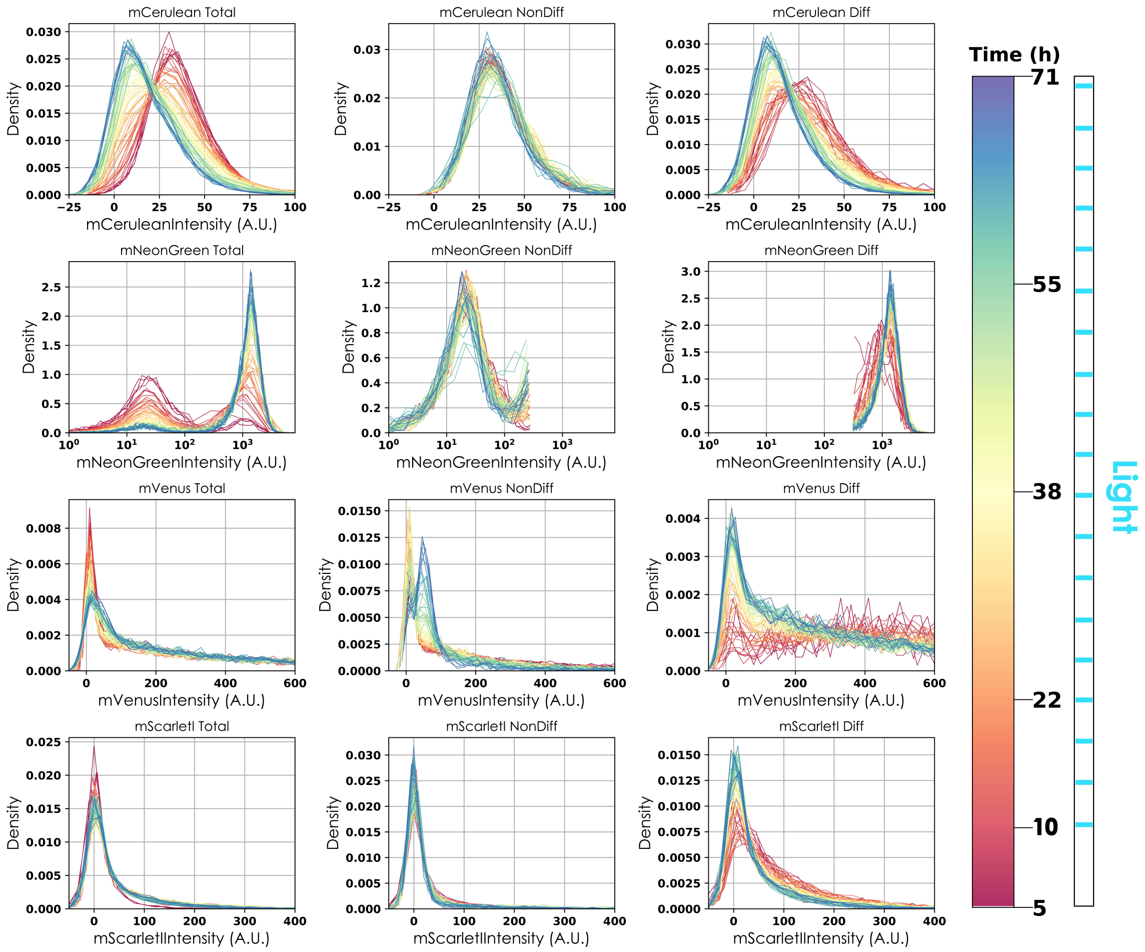


Figure S.13: **Distribution dynamics of the four fluorescent reporter proteins for the experiment with 2min of light every 4h**. Color coding indicates the time of the experiment. Rows correspond to the different fluorescent reporters, columns to different sub-populations (left: all cells, middle: undifferentiated cells, right: differentiated cells) after classification based on mNeonGreen abundance. We note that due to changing sizes of sub-populations, fluorescence distributions need to be extracted from varying (sometimes low) cell numbers.

### S.7 Experimental methods and data analysis

#### S.7.1 Experimental methods

**Cloning.** Plasmids used in the study were generated via the modular cloning approach, Golden-Gate cloning. Yeast tool kit [11] was used to assemble backbones used in the study and served as a library of standard parts (promoters, terminators and connectors). Additional parts were generated either via DNA synthesis or Phusion PCR in the laboratory. Thermocompetent E. coli cells were transformed with the Golden Gate reaction mixture using heat shock transformation. All plasmids were isolated using standard miniprep kits (Macheray & Nagel, and Qiagen). Sequences of plasmids can be found in Supplementary table 1.

**Yeast strains.** All strains used in this study are derived from BY4741 [MATa his3Δ1 leu2Δ0 met15Δ0 ura3Δ0]. Cells were transformed with linearized integrative plasmids or circular 2-micron plasmids using standard Lithium Acetate transformation. Standard auxotrophic markers Uracil and Leucine were used for selection. Integrative plasmids carrying LEU2 and URA3 targeted their endogenous loci. Selection was carried out on agar plates made from standard defined media (Sigma Aldrich Yeast Nitrogen Base) containing 2% glucose and devoid of the respective auxotrophic nutrient (Sigma Aldrich Uracil, and Leucine drop-out media supplements) during 2 to 3 days at 30°C in the dark. Strains used in this study and their genotypes are summarised in Supplementary table 1. Glycerol stocks were made with overnight cultures in selective media when the plasmid was not chromosomally integrated. Such strains were revived on selective plates. On the other hand, strains harboring chromosomally integrated plasmids were stocked from overnight cultures grown in YPD and were revived on YPD agar plates. All stocks were covered in aluminium foil prior to storage. Light sensitive strains were grown in the dark. All manipulations were performed in the presence of red light.

**Growth and culturing conditions.** Strains carrying integrations were grown in non-selective media (Formedium LoFlo Yeast Nitrogen Base supplemented with complete supplement mixture and 2% glucose (SC media)) for all experiments except colony counting assays. Strains harboring 2-micron plasmids were grown in selective media (Formedium LoFlo Yeast Nitrogen Base supplemented with Sigma Aldrich Uracil drop-out supplement and 2% glucose (SD URA-)) for all experiments except plasmid loss assays. Overnight (ON) cultures were started by picking a single colony from a freshly streaked plate in 50 ml Falcon tubes shaking at 200 rpm at 30°C. ON cultures were diluted 1:50 to start precultures on the day of the experiment in 50 ml Falcon tubes shaking at 200 rpm at 30°C. Cultures were allowed to grow for at least 3 hours before loading in individual reactors in the turbidostat platform.

**Turbidostat platform.** Cultures were maintained in exponential phase and monitored at the single-cell and population levels using our previously published custom continuous culture platform equipped with LEDs and capable of controlling the cell density (Figure S.14) [12]. Cultures were grown in the dark until they reached exponential phase and light induction was started only after the growth rate stabilized. Induction was carried out by LED strips (Adafruit NeoPixel Digital RGB LED Strip). An intensity of 40 was used for all experiments (out of a maximum of 255). Sampling from the turbidostat was automated and programmable. Sampled cultures were diluted 20 times with the help of a pipetting robot and passed through the cytometer. The reactor (culture vessel including pumps, tubing, and filters) was autoclaved before each experiment. The experiments followed a grow and dilute program where cultures could grow until an optical density (OD) of 0.6 and were then diluted back to OD 0.4. Information pertaining to dilutions was stored, in addition to the OD and LED status, as csv files. The slope of a linear curve fit to the log of OD data with time was used to estimate the growth rate. Subsequent analysis of the data was done in Python and MATLAB.

**Cytometry and data analysis.** A Guava EasyCyte BGV 14HT benchtop cytometer was used to acquire all cytometry data shown here with constant settings and no compensation during acquisition. 5000 events were recorded at each timepoint in every experiment. Gating was done using kernel density based methods and deconvolution was performed on acquired cytometry data using a linear algebra approach as described in Bertaux et al. [12]. Nonetheless, a summary is provided in Section S.7.2, Figure S.14 and Figure S.16. To calculate the differentiated fraction, a threshold of 300 (a.u.) was applied on deconvolved FSC normalized mNeonGreen fluorescence after gating and doublet removal (Section S.7.2, Figure S.14 and Figure S.18). Cells above the differentiation threshold were classified as differentiated. Python and MATLAB were used for data analysis and visualization.

**Colony counting.** ON cultures were started by picking 3 colonies for both the integrated and 2-micron version of our system on the day before the experiment in 50ml Falcon tubes shaking at 200 rpm at 30°C in SD URA-. ON cultures were diluted to 10ml at OD 0.1 on the morning of the experiment in 50ml flat bottom Erlenmeyer flasks for 6 hours to ensure growth in exponential phase. At the end of 6h, serial dilutions were made from each culture by adding selective media and $200\mu l$ of each dilution was plated on SD URA- and SC agar plates for all 3 replicates. 10 glass beads were added to both SC and SD URA- plates and both the plates were agitated at the same time retracing an imaginary inverted L up and down a total of 8 times [13]. After 48h of growth at 30°C, CFUs were counted manually for all plates.

**Plasmid loss assay.** Cells were grown in selective media overnight in duplicates, ON cultures were started by picking 3 colonies for both the integrated and 2-micron version of the system on the day before the experiment in 50ml Falcon tubes shaking at 200 rpm at 30°C in SD URA-. ON cultures were diluted 1:50 on the morning of the experiment in 50ml Falcon tubes shaking at 200 rpm at 30°C in SD URA-for 6 hours before loading in the bioreactor. The loading in the bioreactor led to a further 3-fold dilution of the preculture but this time in SC media. Cytometry measurements were immediately started upon loading using the automated cytometry functionality of the platform.

#### S.7.2 Data generation and analysis.

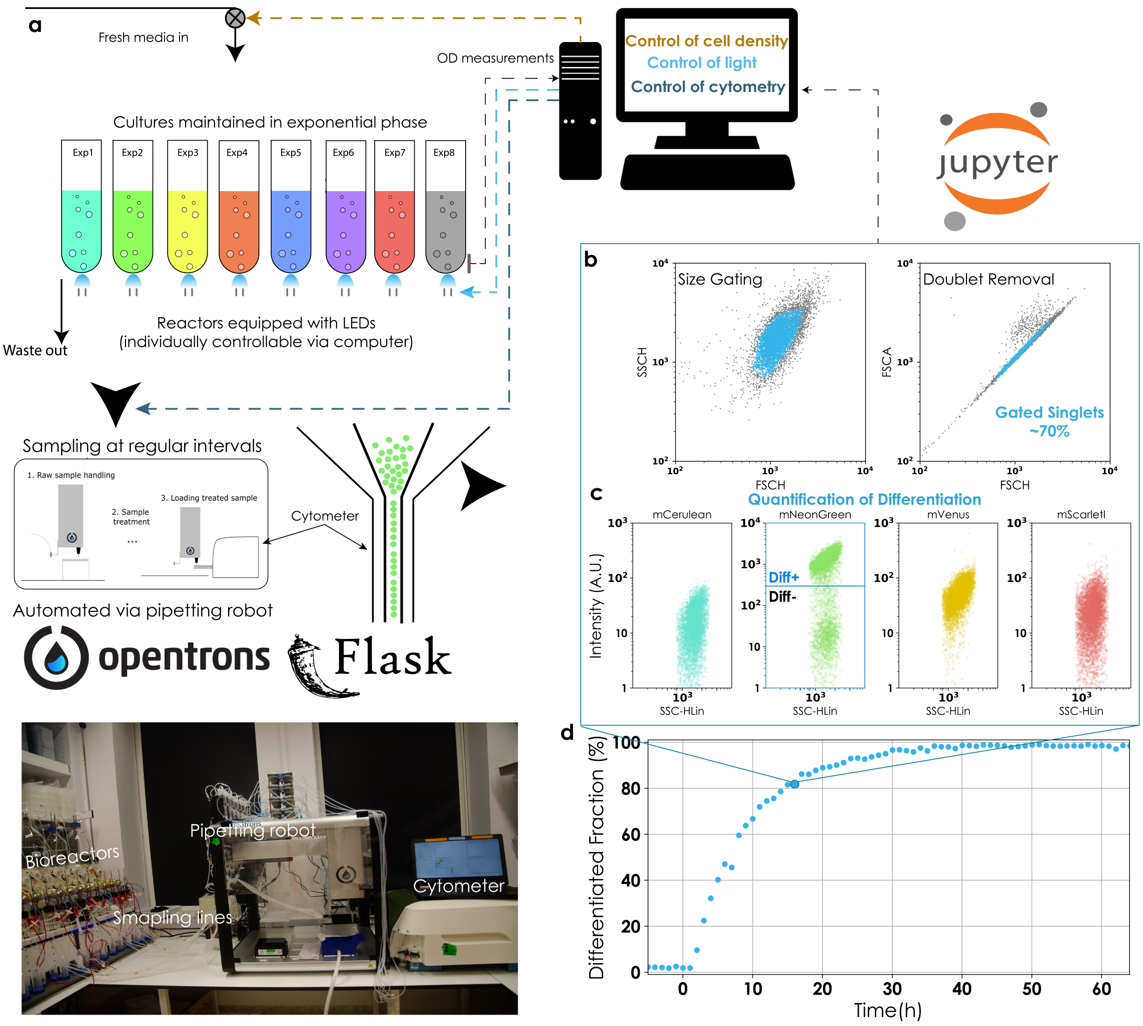


Figure S.14: **Experimental platform and data analysis overview.** **(a)** Up to 8 experiments can be conducted in parallel with regular cytometry measurements and individual OD and LED induction control. Measurements with the cytometer are automated via a pipetting robot that is controlled with the Flask app. Raw fcs files are parsed for each experiment at every timepoint and stored as a single csv file. An image of the physical setup of the experimental platform is shown (bottom left). The software to control and operate the turbidostat platform is developed in Python and implemented with the help of Jupyter notebooks. **(b,c)** Size gating and doublet removal (b) and quantification of differentiation (c) for representative data from a single timepoint (enlarged circle in panel d) for the integrated version of the system induced with continuous light. The FSCH vs SSCH scatterplot is shown as an indicator of cell size and presence of doublets can be inferred from a FSCA vs. FSCH scatterplot. FSC normalized mNeonGreen fluorescence is used to classify cells as differentiated if they exceed the threshold value of 300 (a.u.). Circles represent individual cells. **(d)** Differentiation dynamics over time for a continuously induced culture. Circles represent the fraction of differentiated cells in the population. At the representative time point (large circle), two sub-populations exist in the culture. Light-induction was started at $t = 0$.

We used our previously described turbidostat platform [12] to continuously culture cells and conduct time course experiments. Briefly, the platform allows us to monitor 16 cultures in parallel with regular OD measurements and maintain them at a target cell density. The OD measurements and dilution data were used to estimate the growth rate. Culture vessels are equipped with LEDs and can be stimulated independently with light. Samples from the vessels are collected in a 96-well plate and, with the help of a pipetting robot, loaded in the cytometer. The cytometer acquisition is controlled with the help of click and point software. Figure S.14 gives a general overview of the experimental platform and data analysis.

All cytometry measurements were performed with a Guava EasyCyte BGV 14HT benchtop flow cytometer. Settings and gains were kept constant for all the experiments. 5000 events were acquired for each sample without any compensation. Dilutions were done with a pipetting robot such that the cell density was kept between 200 (to have 5000 events in the acquisition window) and 600 cells/$\mu l$ (to ensure $> 90\%$ singlets). Size gating and doublet removal were done using kernel density based methods. Singlets were selected based on deviation from linearity in Forward Scatter Height (FSC-H) vs. Forward Scatter Area (FSC-A). Cells were scored and a threshold was defined above which cells were classified as doublets and removed from analysis. For size gating, 2D kernel density estimates were obtained using the SciPy Gaussian kde package on Forward Scatter (FSC-H) vs. Side Scatter (SSC-H) and regions of density lower than a threshold were removed (Figure S.14b). The two thresholds were kept constant for all measurements.

Due to significant overlap of fluorescence spectra, it is difficult to observe all four reporter proteins simultaneously (Figure S.15a). To mitigate this, we implemented a deconvolution approach previously described in Bertaux et al. [12]. Briefly, 4 single fluorescent protein control strains (mCerulean, mNeonGreen, mVenus, mScarlet-I) with the same promoter and terminator, and integrated in the same locus, were used to determine the spectral signature of each fluorescent protein across the 12 channels of the cytometer. These signatures were then used in a linear algebra framework to calculate the individual fluorescence of each fluorophore in a strain harboring all 4 of the fluorescent proteins. We note that prior to deconvolution, autofluorescence in each channel was subtracted from the raw cytometry data. Autofluorescence was estimated by culturing a wildtype (WT) strain (BY4741) complemented with a URA3 cassette that expresses no fluorescent protein. Fluorescence distributions for single-color control and the 4-color strain were in good agreement after deconvolution (Figure S.15b).

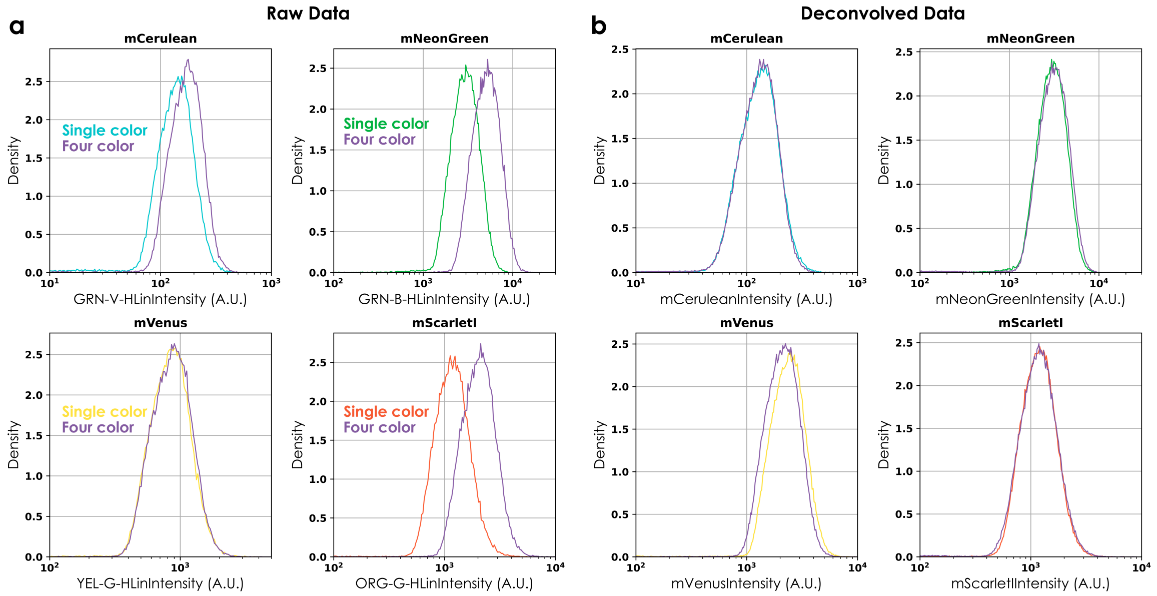


Figure S.15: Fluorescence distributions before (a) and after (b) deconvolution of the 4-color strain are compared to fluorescence distributions of single-color strains. For each fluorescent protein, the signal in the most distinguishing channel is plotted in (a), as noted on the x-axes of sub-panels. Cells were size gated and doublets were removed. Histograms are composed of $> 100,000$ cells.

There was a slight disagreement between the single-color and 4-color deconvolved fluorescence for mVenus (5% shift in median value). This is perhaps due to errors in deconvolution but might also stem from epigenetic differences in the gene expression loci for single color (URA3 locus) and 4 color (HO locus) strains [14].

Figure S.16a and Figure S.17a show raw fluorescence values in the most distinguishing channel for each of the four fluorescent proteins present in the 4-color strain in the dark (grey), when 99% of the population is undifferentiated, and in continued presence of light (blue) after 99% of the population has differentiated. Prior to deconvolution, the raw data possessed several characteristics suboptimal for analysis. Notably, due to low fluorescence of mCerulean, the undifferentiated population (possessing mCerulean) overlaps significantly with the differentiated population (possessing no mCerulean) in the GRN-V channel. Seemingly, undifferentiated cells possess 5-6 times higher fluorescence than the WT strain in the GRN-B channel. Furthermore, the fluorescence in the YEL-G channel, misleadingly, suggests large differences in mVenus fluorescence between differentiated and undifferentiated cells. Lastly, observing the ORG-G channel, we find that cells in the dark possess 3-4 fold higher fluorescence compared to the WT strain prompting the misleading conclusion that EL222 is activated without induction. The situation becomes significantly worse for the plasmid-based system such that it becomes impossible to observe two well-separated populations for differentiated and undifferentiated cells (Figure S.17a, top right). Furthermore, it appears from YEL-G fluorescence that cells stop losing the plasmid upon differentiation (Figure S.17a, bottom left).

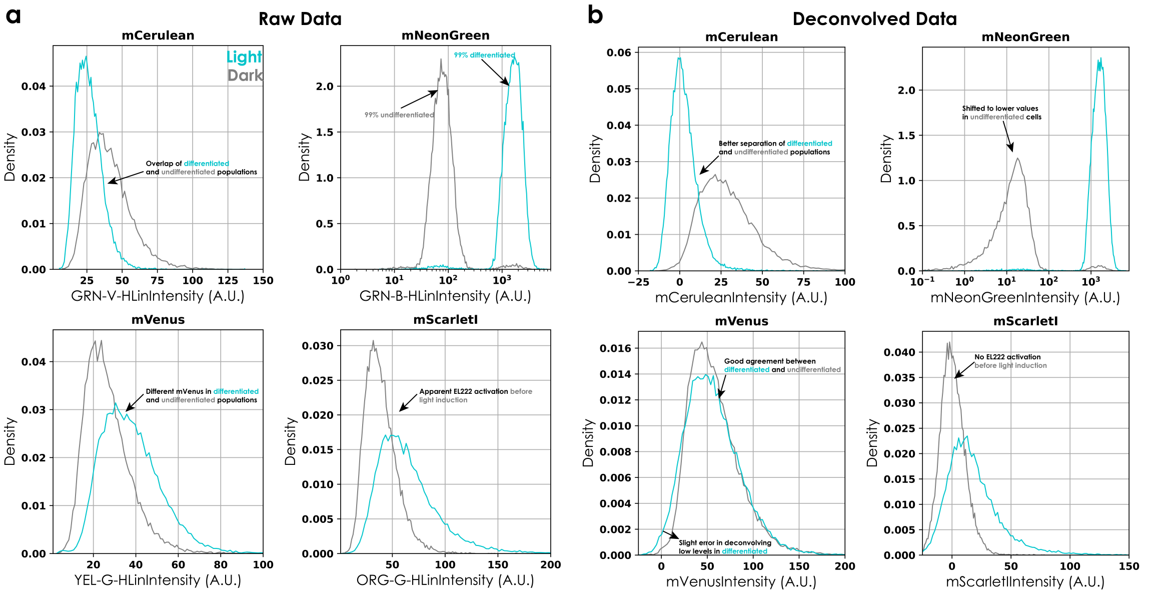


Figure S.16: Fluorescence distributions before (a) and after (b) deconvolution of the 4-color strain for the integrated system in light (blue) and dark (grey). Data is represented as in Figure S.15. Cells were size gated and doublets were removed. Histograms are composed of $> 10,000$ cells.

Deploying our deconvolution strategy allows us to deconvolve raw fluorescence in 4-color strains. After deconvolution, all the above-mentioned incongruities either disappear completely or are significantly diminished. We note that devonvolution of low levels of mVenus in the presence of mNeonGreen (differentiated cells) yields noisy estimates (Figure S.16b, bottom left) that lead to a broad spread around zero for the plasmid strain (Figure S.17b, bottom left).

For the quantification of fluorescence, we normalized single-cell fluorescence values after deconvolution by the cells' FSC values to reduce effects of cell size variability. To preserve units, values were scaled by multiplying with the mean FSC of the population. We found that normalizing by FSC led to tighter distributions, except for mScarlet-I fluorescence, perhaps due to low fluorescence values. Moreover, we found a persistent population of cells that possessed neither mNeonGreen nor mCerulean fluorescence at detectable levels. Since our biological circuit precludes their existence, these spurious cells were filtered out by applying thresholds on mCerulean (10a.u.) and mNeonGreen fluorescence (300a.u.) and excluding cells that have neither mCerulean nor mNeonGreen fluorescence above the respective threshold. Typically these cells made up less than 0.1% of the total cells. The existence of these cells could be a consequence of sampling in open plates for cytometry or contamination in the sampling lines.

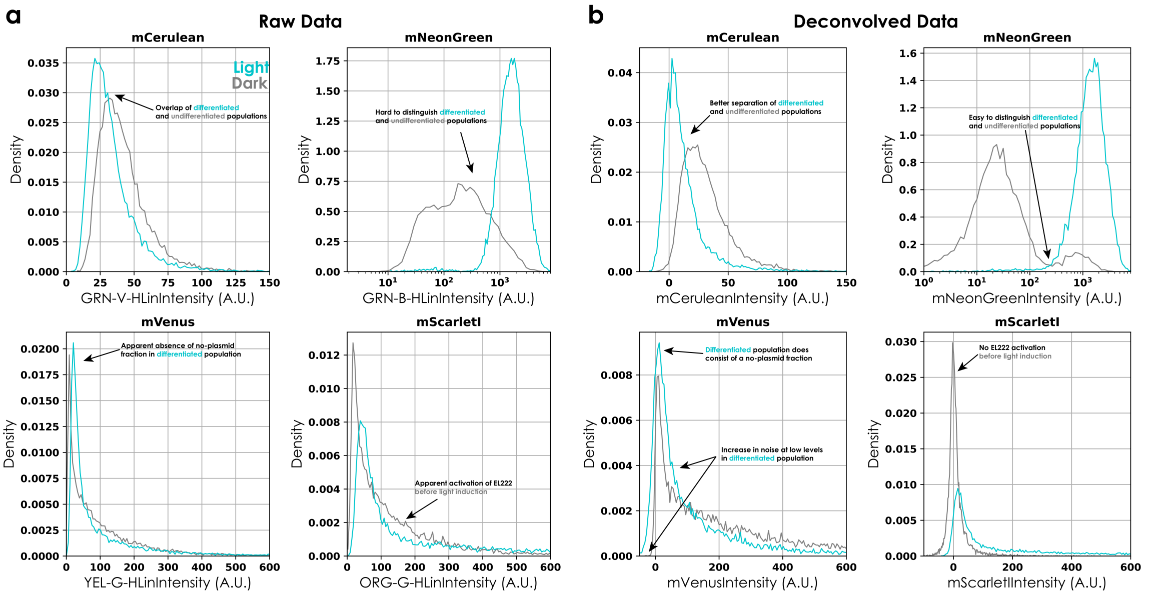


Figure S.17: Fluorescence distributions before (a) and after (b) deconvolution of the 4-color strain for the plasmid-based system in light (blue) and dark (grey). Data is represented as in Figure S.15. Cells were size gated and doublets were removed. Histograms are composed of $> 10,000$ cells.

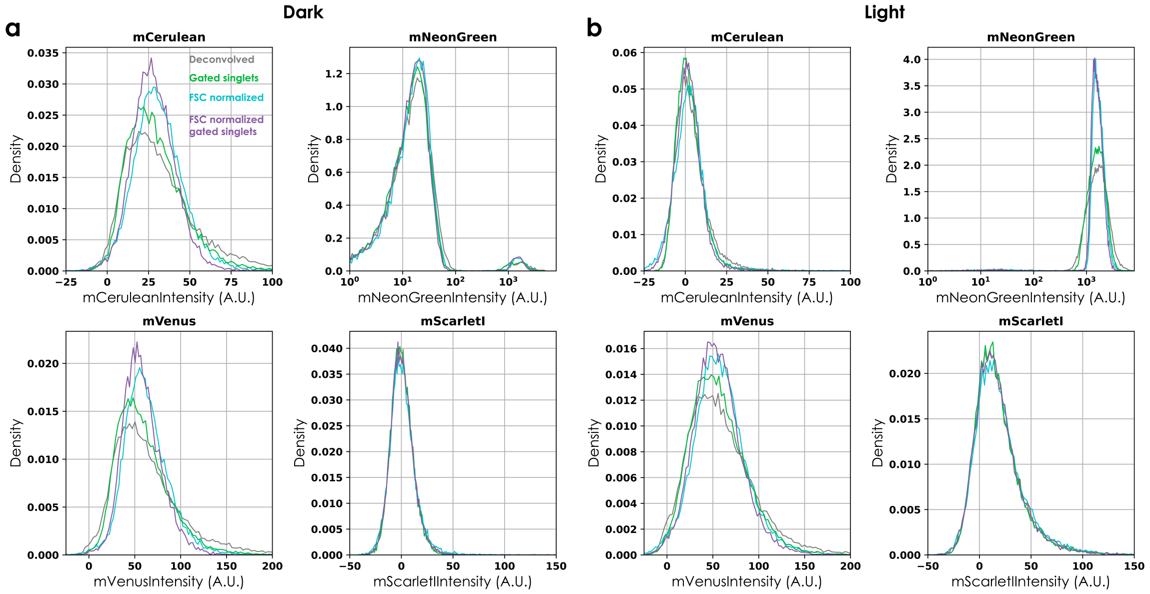


Figure S.18: Fluorescence distributions of the four reporter proteins of the 4-color strain for the integrated system in dark (a) and light (b) for deconvolved (grey), gated singlets (green), FSC normalized deconvolved (blue), and FSC normalized gated singlets (purple) data. Histograms are composed of $> 10,000$ cells.

### References


[1] Friedman N, Cai L, Xie S (2006) Linking stochastic dynamics to population distribution: An analytical framework of gene expression. *Physical Review Letters* 97:168302.

[2] Benzinger D, Khammash M (2018) Pulsatile inputs achieve tunable attenuation of gene expression variability and graded multi-gene regulation. *Nature Communications* 9:3521.

[3] Shahrezaei V, Swain P (2008) Analytical distributions for stochastic gene expression. *Proceedings of the National Academy of Sciences of the USA* 105:17256–17261.

[4] Lunz D, Batt G, Ruess J, Bonnans J (2021) Beyond the chemical master equation: stochastic chemical kinetics coupled with auxiliary processes. *PLoS Computational Biology* 17:e1009214.

[5] Munsky B, Khammash M (2006) The finite state projection algorithm for the solution of the chemical master equation. *The Journal of Chemical Physics* 124:044104.

[6] Darroch J, Seneta E (1967) On quasi-stationary distributions in absorbing continuous-time finite markov chains. *Journal of Applied Probability* 4:192–196.

[7] Karim A, Curran K, Alper H (2013) Characterization of plasmid burden and copy number in saccharomyces cerevisiae for optimization of metabolic engineering applications. *FEMS yeast research* 13:107–116.

[8] Fang F, et al. (2011) A vector set for systematic metabolic engineering in saccharomyces cerevisiae. *Yeast* 28:123–136.

[9] Gnügge R, Liphardt T, Rudolf F (2016) A shuttle vector series for precise genetic engineering of saccharomyces cerevisiae. *Yeast* 33:83–98.

[10] Sidje RB (1998) Expokit: a software package for computing matrix exponentials. *ACM Transactions on Mathematical Software (TOMS)* 24:130–156.

[11] Lee M, DeLoache W, Cervantes B, Dueber J (2015) A highly characterized yeast toolkit for modular, multipart assembly. *ACS synthetic biology* 4:975–986.

[12] Bertaux F, et al. (2020) Enhancing bioreactor arrays for automated measurements and reactive control with reacsight. *bioRxiv*.

[13] Prusokas A, Hawkins M, Nieduszynski C, Retkute R (2021) Effectiveness of glass beads for plating cell cultures. *Physical Review E* 103:052410.

[14] Bai Flagfeldt D, Siewers V, Huang L, Nielsen J (2009) Characterization of chromosomal integration sites for heterologous gene expression in saccharomyces cerevisiae. *Yeast* 26:545–551.
